# Clonal evolution of hematopoietic stem cells after cancer chemotherapy

**DOI:** 10.1101/2024.05.23.595594

**Authors:** Hidetaka Uryu, Koichi Saeki, Hiroshi Haeno, Chiraag Deepak Kapadia, Ken Furudate, Jyoti Nangalia, Michael Spencer Chapman, Li Zhao, Joanne I. Hsu, Chong Zhao, Shujuan Chen, Tomoyuki Tanaka, Zongrui Li, Hui Yang, Courtney DiNardo, Naval Daver, Naveen Pemmaraju, Nitin Jain, Farhad Ravandi, Jianhua Zhang, Xingzhi Song, Erika Thompson, Hongli Tang, Latasha Little, Curtis Gumbs, Robert Z. Orlowski, Muzaffar Qazilbash, Kapil Bhalla, Simona Colla, Hagop Kantarjian, Rashmi Kanagal Shamanna, Carlos Bueso- Ramos, Daisuke Nakada, P. Andrew Futreal, Elizabeth Shpall, Margaret Goodell, Guillermo Garcia-Manero, Koichi Takahashi

**Author notes:** **Correspondence**: Koichi Takahashi, MD, PhD., Department of Leukemia, Unit 428, The University of Texas MD Anderson Cancer Center 1515 Holcombe Boulevard, Houston, TX 77030, USA.

## Abstract

Normal hematopoietic stem and progenitor cells (HSPCs) inherently accumulate somatic mutations and lose clonal diversity with age, processes implicated in the development of myeloid malignancies^1^. The impact of exogenous stressors, such as cancer chemotherapies, on the genomic integrity and clonal dynamics of normal HSPCs is not well defined. We conducted whole-genome sequencing on 1,032 single-cell-derived HSPC colonies from 10 patients with multiple myeloma (MM), who had undergone various chemotherapy regimens. Our findings reveal that melphalan treatment distinctly increases mutational burden with a unique mutation signature, whereas other MM chemotherapies do not significantly affect the normal mutation rate of HSPCs. Among these therapy-induced mutations were several oncogenic drivers such as *TET2* and *PPM1D*. Phylogenetic analysis showed a clonal architecture in post-treatment HSPCs characterized by extensive convergent evolution of mutations in genes such as *TP53* and *PPM1D*. Consequently, the clonal diversity and structure of post-treatment HSPCs mirror those observed in normal elderly individuals, suggesting an accelerated clonal aging due to chemotherapy. Furthermore, analysis of matched therapy-related myeloid neoplasm (t-MN) samples, which occurred 1-8 years later, enabled us to trace the clonal origin of t-MNs to a single HSPC clone among a group of clones with competing malignant potential, indicating the critical role of secondary mutations in dictating clonal dominance and malignant transformation. Our findings suggest that cancer chemotherapy promotes an oligoclonal architecture with multiple HSPC clones possessing competing leukemic potentials, setting the stage for the selective emergence of a singular clone that evolves into t-MNs after acquiring secondary mutations. These results underscore the importance of further systematic research to elucidate the long-term hematological consequences of cancer chemotherapy.

## Main

The acquisition of somatic DNA mutations is a hallmark of aging and contributes to the development of various age-associated human diseases, including cancer^2^. In hematopoietic stem cells (HSCs), mutation acquisition follows a remarkably linear trajectory over time, with individual HSCs acquiring approximately 16 to 25 mutations per year^1,3–5^. Certain somatic mutations can confer increased fitness to cells, leading to their positive selection (i.e., driver mutations). This results in clonal expansion of cells harboring these driver mutations, a process known as clonal hematopoiesis (CH)^6,7^. The progression of CH contributes to the age-related decline of clonal diversity in HSCs and ultimately to the development of hematologic malignancies ^1,8^.

External stressors or carcinogens can modulate mutation rates and cellular dynamics in human tissues. Factors such as ultraviolet (UV) light, tobacco smoking, and alcohol intake increase somatic mutations in tissues like skin, lung, and esophagus^9–13^. Furthermore, these extrinsic factors alter the fitness landscape of cellular ecosystems, enabling the expansion of cells with mutations that confer a fitness advantage under such conditions^9,12–14^.

The hematopoietic system is also exposed to various external stressors, including cytotoxic chemotherapy, which in rare instances can lead to the development of therapy-related myeloid neoplasms (t-MNs)^15^. DNA sequencing studies of cancer patients previously treated with chemotherapy revealed distinct mutational profiles of CH compared to the general population, with enrichment of mutations in DNA-damage response (DDR) pathway genes, such as *TP53*, *PPM1D*, and *CHEK2*^16,17^. In animal models, DNA-damaging chemotherapy has been demonstrated to promote the selective expansion of HSCs with *TP53* and *PPM1D* mutations^18–20^.

While these previous studies have revealed the relationship between chemotherapy exposure and clonal expansion of specific driver mutations, how chemotherapy treatment influences the overall population dynamics and genomes of individual human HSCs remains unclear. Moreover, how these effects contribute to the development of treatment-induced malignancy is not fully understood. Here, we employed a single-cell colony whole-genome sequencing approach to investigate the effects of chemotherapy on the genome and population dynamics of normal HSCs using peripheral blood stem cells (PBSCs) collected from patients previously treated with chemotherapy. By analyzing the matched t-MN genome, we also elucidated the evolutionary history of t-MN development arising from normal HSCs.

## Results

### Whole-genome sequencing of single-HSPC colonies from patients previously treated with chemotherapy

To study the impact of chemotherapy on the genome of hematopoietic stem and progenitor cells (HSPCs), we analyzed mobilized PBSCs collected from 10 patients with multiple myeloma (MM) ages between 46-65 (**Figure 1a**). These patients were previously treated with various induction chemotherapies and underwent PBSC collection prior to autologous stem cell transplantation (ASCT). Two patients underwent ASCT twice with a different PBSC collection with an interval of 3 years and 15 years, respectively, and PBSCs collected from both time points were studied. **Table 1** describes the detailed clinical characteristics of the 10 patients. Since PBSCs are collected prior to ASCT, cells are exposed to chemotherapies used for induction and mobilization (if chemotherapy is used), but not to the chemotherapies used for transplant conditioning because PBSCs are infused after completing conditioning chemotherapy (**Extended Figure 1a**). For PBSCs collected at the 2^nd^ ASCT, chemotherapy exposure also includes maintenance or salvage therapies given after the 1^st^ ASCT and mobilization therapy prior to the 2^nd^ PBSC collection, in addition to the exposure for the 1^st^ ASCT (**Extended Figure 1b**). Of note, in both scenarios, PBSCs are not exposed to the high-dose chemotherapies used for transplant conditioning. With these factors in mind, the list of prior therapy exposures for HSPCs included melphalan (N = 2), cyclophosphamide (N = 3), doxorubicin (N = 2), vincristine (N = 2), lenalidomide or thalidomide (N = 7), bortezomib (N = 5), radiation (N = 2), and interferon alpha (N = 1) (**Figure 1b**). This variation in the prior therapy allowed us to study the effect of both conventional cytotoxic chemotherapies (melphalan, cyclophosphamide, doxorubicin, and vincristine) and non-cytotoxic therapies (lenalidomide, thalidomide, and bortezomib) on HSPC genomes.

**Figure 1.**
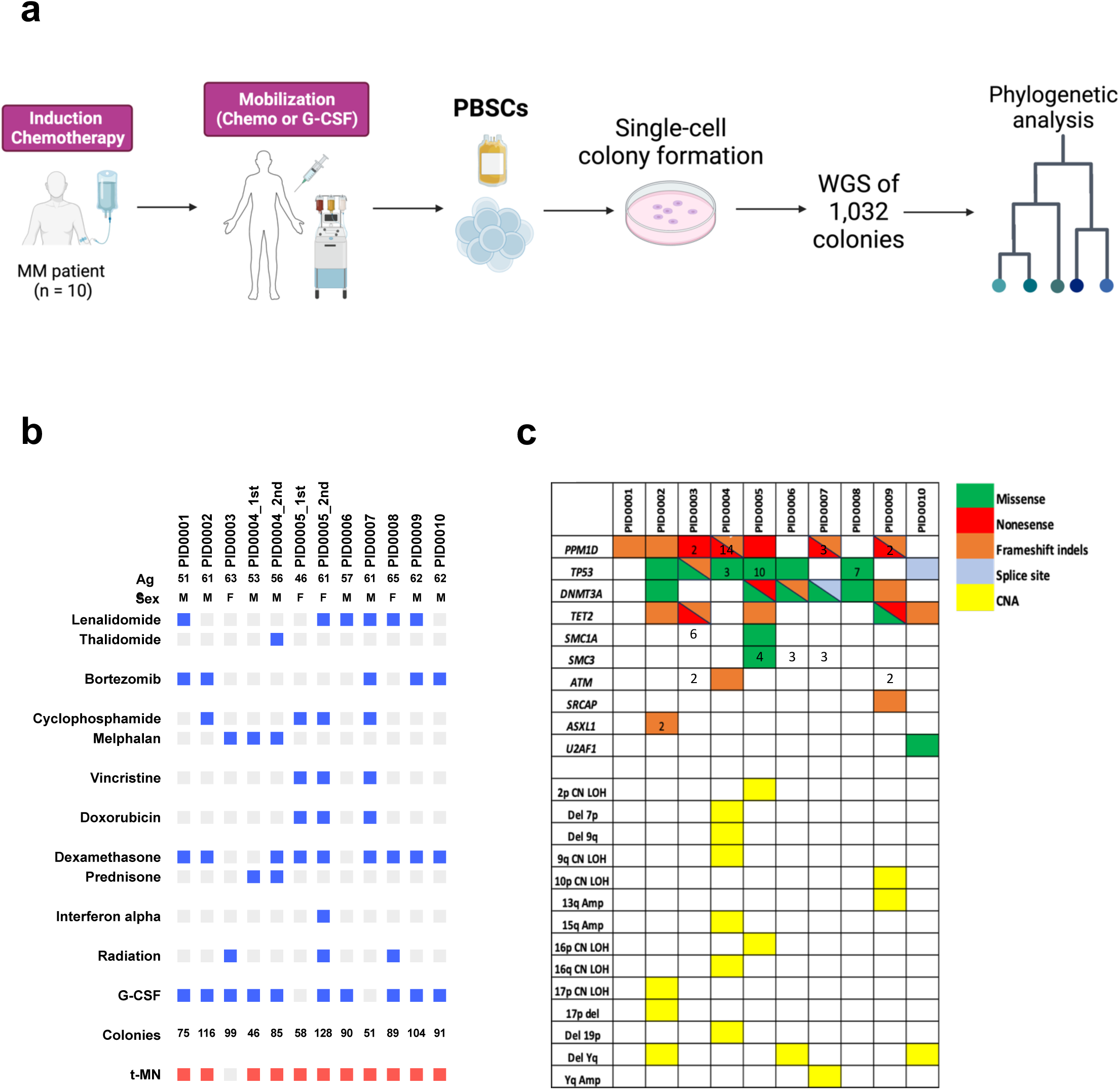
Overview of study design and cohort. (**a**) Diagram illustrating the experimental workflow. (**b**) Figure summarizing patient demographics, specific chemotherapeutic agents encountered by the analyzed hematopoietic stem and progenitor cells (HSPCs), the quantity of colonies evaluated, and the occurrence of therapy-related myeloid neoplasms (t-MN). Bright red boxes denote t-MN samples subjected to whole genome sequencing, while pink boxes indicates t-MN samples that were unavailable for sequencing analysis. (c) Oncoplot of driver mutations and copy number alterations (CNAs) identified in at least one colony per patient. The numeric value within each box represents the count of distinct mutations identified within the same gene. MM = multiple myeloma, PBSCs = peripheral blood stem cells, WGS = whole genome sequencing.

**Table 1.** Clinical characteristics of 10 patients with multiple myeloma (MM)

| Patient ID | Sex | Induction Chemotherapy | Mobilization | Conditioning | Age at 1<sup>st</sup> PBSC collection | Chemotherapy after 1<sup>st</sup> auto | Mobilization for 2<sup>nd</sup> auto | Conditioning for 2<sup>nd</sup> auto | Age at 2<sup>nd</sup> PBSC collection | # of colonies (1<sup>st</sup> / 2<sup>nd</sup>) | t-MNs (Interval from last auto) |
| --- | --- | --- | --- | --- | --- | --- | --- | --- | --- | --- | --- |
| PID0001 | Male | Lenalidomide/Bortezomib/Dexamethasone | G-CSF | Melphalan | 51 |  |  |  | - | 75 | t-MDS (3y) |
| PID0002 | Male | Cyclophosphamide/Bortezomib/Dexamethasone | G-CSF | Melphalan | 61 |  |  |  | - | 116 | t-MDS (1y) |
| PID0003 | Female | Melphalan + XRT | G-CSF | Melphalan + Lenalidomide | 63 |  |  |  | - | 99 | - |
| PID0004 | Male | Melphalan + Prednisone | G-CSF | Topotecan, Cyclophosphamide, Melphalan | 53 | Thalidomide, Dexamethasone | G-CSF | Melphalan | 56 | 133 (46/87) | t-MDS (4y) |
| PID0005 | Female | Vincristine, Doxorubicin, Dexamethasone (VAD) | Cyclophosphamide | Thiotepa, Busulfan, Cyclophosphamide | 46 | IFN-α, XRT, Lenalidomide | G-CSF | Melphalan, Lenalidomide | 61 | 177 (49/128) | t-MDS (6y) |
| PID0006 | Male | Lenalidomide | G-CSF | Melphalan | 57 |  |  |  | - | 90 | t-MDS (7y) |
| PID0007 | Male | Lenalidomide, Bortezomib, Dexamethasone | CVAD | Melphalan | 61 |  |  |  | - | 52 | t-MDS (1y) |
| PID0008 | Female | Local irradiation, Lenalidomide, Dexamethasone | G-CSF | Melphalan | 65 |  |  |  | - | 89 | t-MDS (6y) |
| PID0009 | Male | Lenalidomide, Bortezomib, Dexamethasone | G-CSF | Melphalan | 62 |  |  |  | - | 104 | t-MDS (1y) |
| PID0010 | Male | Bortezomib, Dexamethasone | G-CSF | Melphalan | 62 |  |  |  | - | 88 | t-MDS (3y) |

**Abbreviations:** G-CSF: Granulocyte colony stimulating factor, CVAD: cyclophosphamide, vincristine, doxorubicin, dexamethasone, XRT: radiation, IFN: interferon, PBSC: peripheral blood stem cells, t-MNs: therapy-related myeloid neoplasms, t-MDS: therapy-related myelodysplastic syndromes

PBSC samples were cultured in semi-liquid methylcellulose media to generate single-cell-derived colonies. Genomic DNA collected from individual single-cell derived colonies was analyzed by whole genome sequencing (WGS). We sequenced a median of 89 colonies per sample (range: 46-128), totaling 1,047 colonies. WGS achieved median 29x coverage (range: 14-68x). Putative somatic mutations were identified in each colony after computationally filtering potential germline variants, sequencing artifacts, and mutations that are likely acquired during *in vitro* culture, using bioinformatic approaches modified from previous studies^4,10^. We analyzed the distribution of variant allele frequency (VAF) of somatic mutations in individual colonies to confirm the single-cell origin, and colonies that do not have VAF peak at 50% or having multiple peaks were removed from further analysis, as they are likely merged colonies (**Extended Figure 2**). Consequently, 1,032 colonies passed the quality control and were analyzed further.

A median of 7 driver mutations (range: 0-19) and 1 copy number alterations (CNAs, range: 0-5) were detected per sample (**Figure 1c**). The most frequently detected driver mutations were *PPM1D* and *TP53* mutations in 7 samples each, followed by *DNMT3A* mutations in 6 samples and *TET2* mutations in 5 samples. CNAs were rare and the only recurrent abnormality was chromosome Yq deletion in 3 samples. The significant enrichment of *PPM1D* and *TP53* mutations, both of which are involved in DNA-damage response (DDR) pathways, was consistent with the high prevalence of these mutations found in therapy-related CH^17^, and in contrast to the mutation profiles in normal individuals^1^.

### Mutation burden and signatures in post-treatment HSPCs

Studies have shown that the mutation rate in normal HSPCs follows a linear trajectory over time, with each single HSPC acquiring approximately 16-25 single-nucleotide variants (SNVs) per year^3,4^. Using this as a benchmark, we plotted the number of somatic SNVs in HSPCs treated with chemotherapy (hereafter, called post-treatment HSPCs) (**Figure 2a**). Despite the history of chemotherapy treatment, the number of somatic SNVs in post-treatment HSPCs from eight out of ten patients followed the mutation rate expected in normal HSPCs. In contrast, three HSPC samples from two patients (one patient had two samples collected at different time points) showed a significant deviation from the age-anticipated mutation burden, demonstrating an increase of approximately twice or more in the number of somatic SNVs (**Figure 2a**, PID0003, and PID0004). Similar trend was also observed in indels and multiple nucleotide variants (MNVs) (**Extended Figure 3**). Notably, both of these two patients had undergone melphalan-containing therapies.

**Figure 2.**
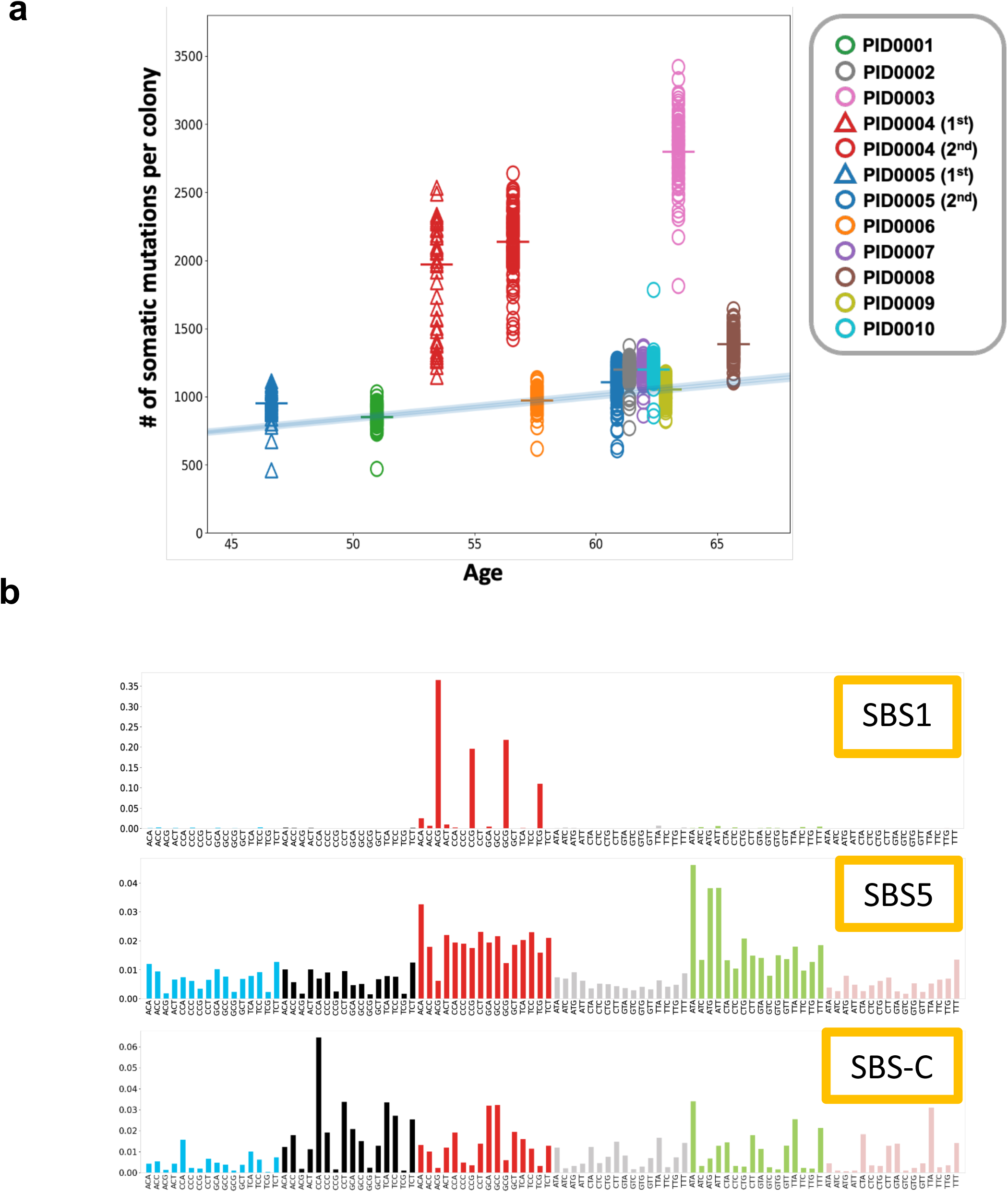
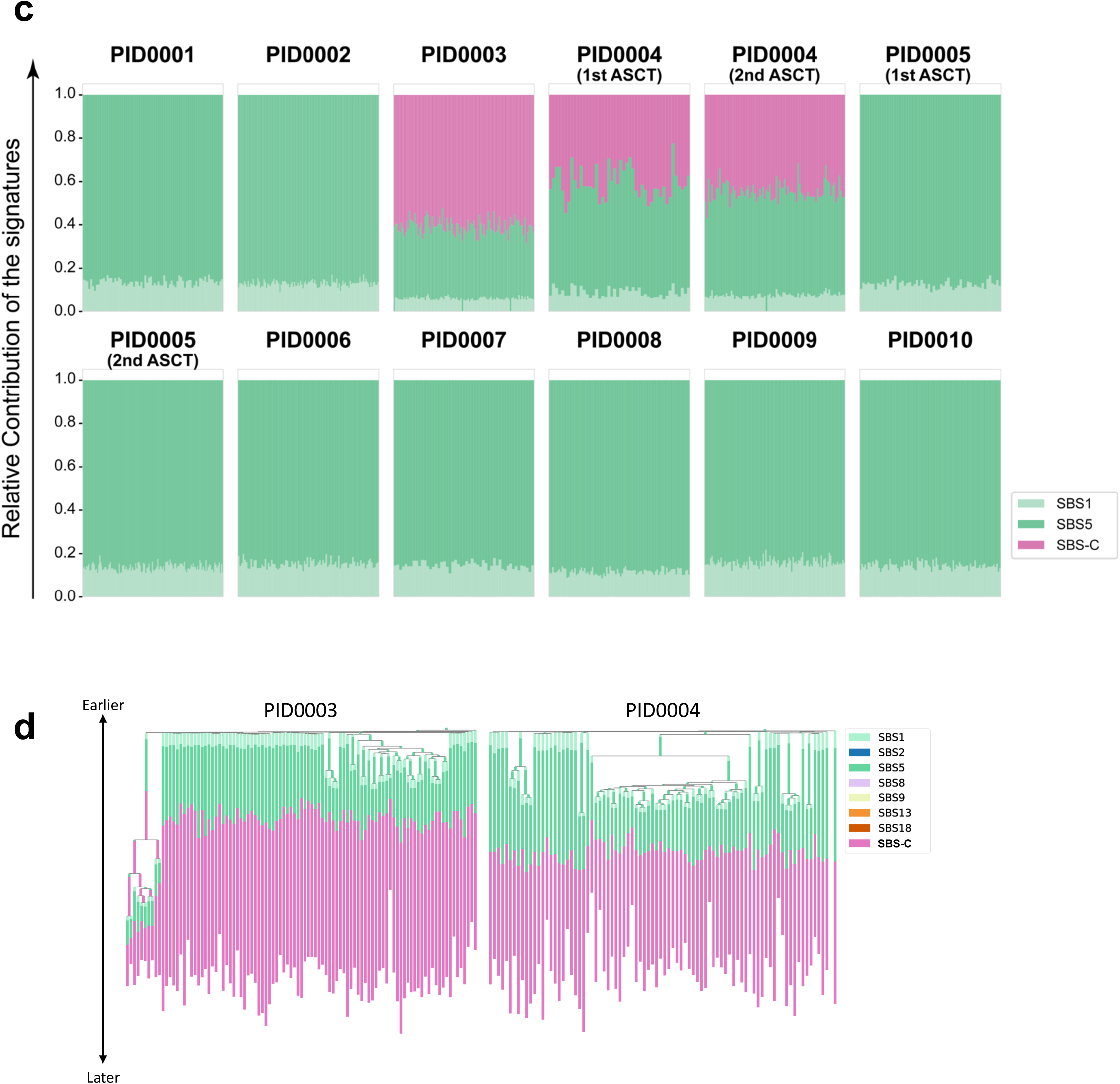
Analysis of somatic mutations and their signatures in HSPC colonies. (**a**) Scatter plot of somatic single nucleotide variants (SNVs) per colony against age, with the blue line representing the expected normal mutation rate as established by Mitchell et al. 2022. (**b**) Three distinct SNV signatures deduced from the sequencing of 1,032 colonies. SBS-C is closely related, with a 90% cosine similarity, to the SBS-MM1 signature identified by Rustad et al. 2020. (**c**) A stacked bar chart illustrating the frequency distribution of SBS-1, SBS-5, and SBS-C signatures across all SNVs in individual colonies. (**d**) Phylogenetic trees for PID0003 and PID0004, incorporating mutation signatures. SBS-C is apparent only in mutations acquired later in the patients’ lives.

In order to determine the causative relationship between melphalan treatment and the observed increase in somatic mutations, we extracted mutation signature from SNVs. In total, 3 mutation signatures were identified (**Figure 2b**). Two were consistent with the known clock-like signatures, SBS1 and SBS5. SBS-C was exclusively found in the two patients exposed to melphalan. This signature showed cosine similarity of 90% with the previously identified signature SBS-MM1, which is a putative melphalan-associated signature detected in myeloma cells^21^ (**Extended Figure 4**). Approximately 60% and 50% of PID0003 and PID0004 mutations, respectively, were attributed to SBS-C, matching the incremental increase of the mutations in these samples compared to others (**Figure 2c**). SBS-C was detected across all colonies with equal proportion, potentially suggesting that melphalan affected HSPCs independent of cell-cycle state, which is consistent with the known mode of action of melphalan^22^. Phylogeny analysis projecting mutation signatures shows that SBS-C is acquired at a later time in PID0003 and PID0004’s life, consistent with the fact that these mutations are acquired after treatment exposure (age of exposure 53 years and 52 years for PID0003 and PID0004, respectively **Figure 2d**). Taken together, these findings provide evidence of a causal link between melphalan exposure and the elevated somatic mutational burden in these HSPCs. None of the other patients’ HSPCs showed treatment-related signatures and only clock-like signatures (SBS1 and SBS5) were observed^23^.

In contrast to melphalan treatment, HSPCs exposed to cyclophosphamide (PID0002, PID0005, and PID0007), an alkylating agent similar to melphalan, did not display an increased number of mutations or treatment-related signatures. This disparity is likely attributed to the known function of HSCs in metabolizing cyclophosphamide into an inactive form via aldehyde dehydrogenase (ALDH), thus evading cyclophosphamide-induced DNA damage, whereas melphalan directly triggers DNA adduct formation in HSCs^22,24,25^. These findings underscore the differential impact of chemotherapy on the HSC genome, and suggest that distinct chemotherapy agents, or even drugs within the same class of agents, may have a variable effect on HSPC genomes *in vivo*.

### Clonal architecture and diversity of post-treatment HSPCs

We then reconstructed phylogenetic trees of the post-treatment HSPCs using shared and unique somatic SNVs of individual colonies. Since treatment-related mutations like SBS-C would confound the clonal relationship and molecular clock, we removed SBS-C-related mutations and only used clock-like signature mutations (SBS1 + SBS5) for phylogeny building (**Figure 3a-b**). The phylogenies were also annotated with known hematologic driver mutations or CNAs to understand the relationship between expanded clades and driver mutations.

**Figure 3.**
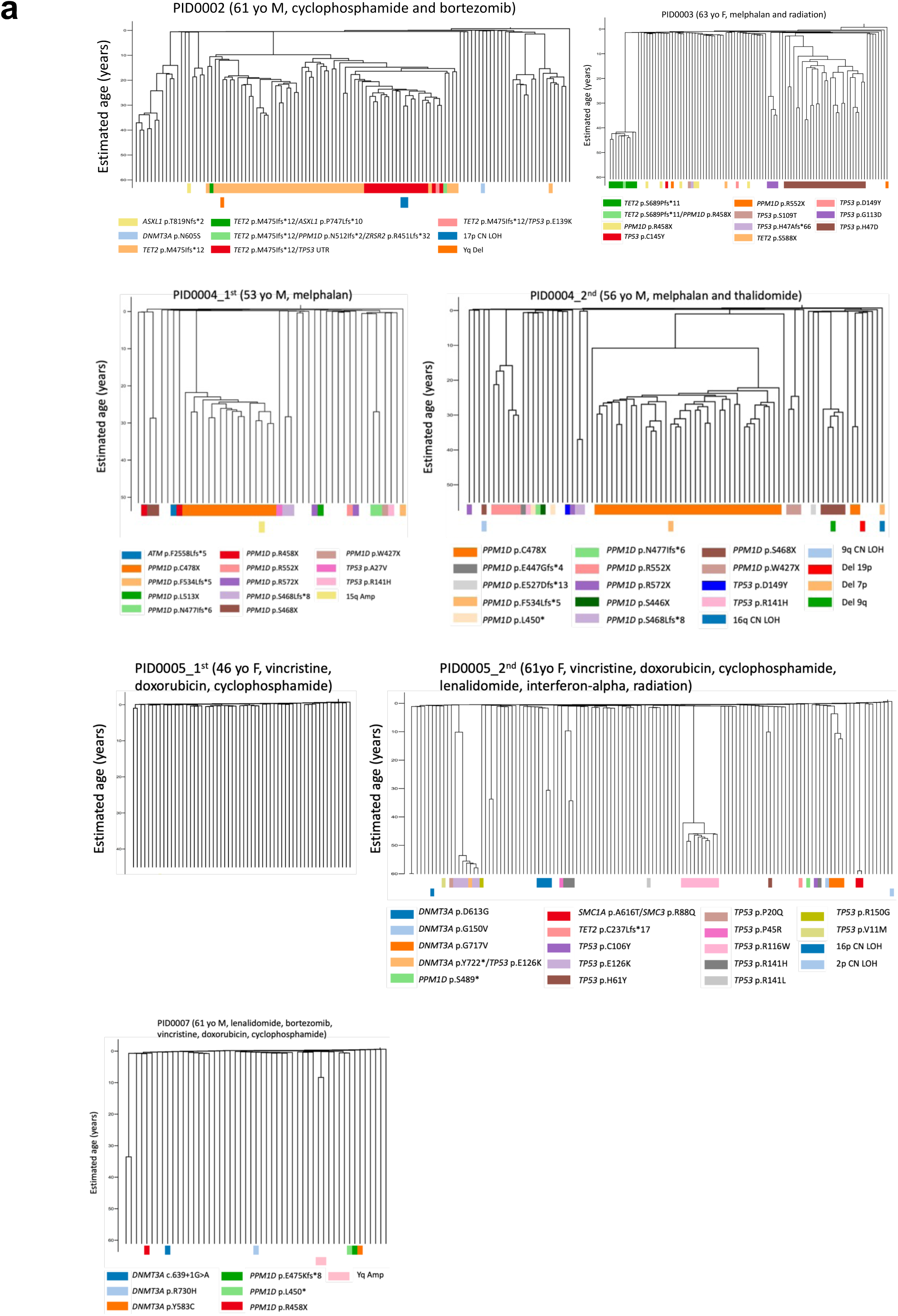
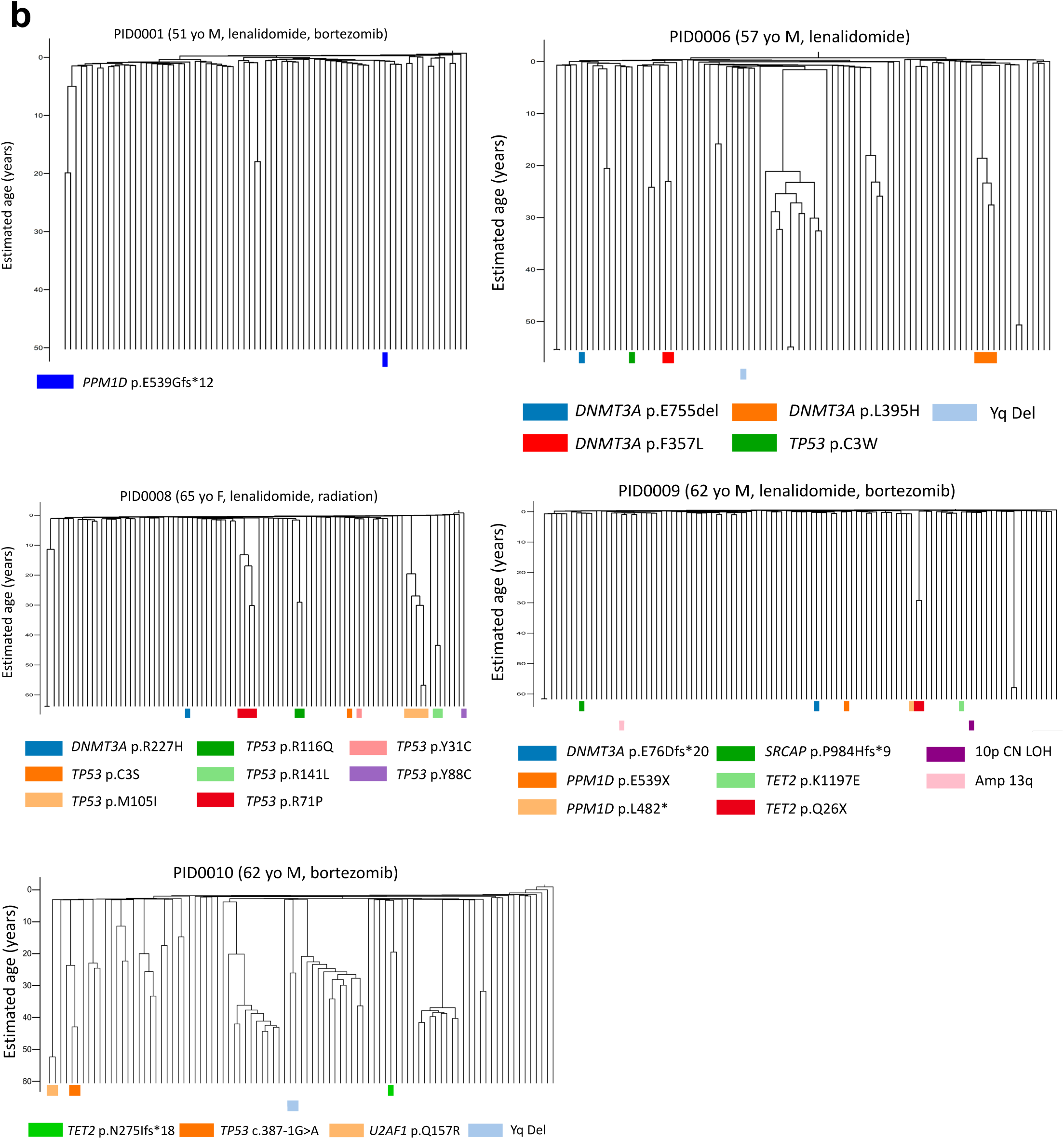
Ultrametric phylogenetic trees constructed from post-chemotherapy HSPCs. These trees are based on SNVs identified in individual colonies, excluding those associated with the SBS-C signature. The trees are further detailed at the bottom, indicating the presence of driver mutations (top row) and copy number alterations (bottom row). (**a**) Trees corresponding to samples that underwent cytotoxic chemotherapy treatments. (**b**) Trees for samples treated with non-cytotoxic chemotherapeutic agents.

The previous study showed that the clonal diversity of HSPCs decreases after age 70 in normal individuals^1^. Qualitatively, some of the samples from the current cohort of post-treatment HSPCs showed pronounced oligoclonality compared to age-matched normal individuals published previously^1^, which is evident from multiple large clades within the same patient (**Figure 3a-b**). Quantitatively, the Shannon diversity index varied significantly among our patients’ samples and a subset of the samples showed a significantly lower diversity index that is comparable to that of normal individuals with age above 75, despite our cohort having patients ages 46-65 (**Figure 4a**). When we compared the diversity index between patients treated with conventional cytotoxic chemotherapies (melphalan, cyclophosphamide, doxorubicin, and vincristine-treated samples) and non-cytotoxic therapies (lenalidomide, thalidomide, and bortezomib-treated samples), HSPCs showed a trend of lower diversity in patients who were treated with cytotoxic chemotherapies compared to the ones received non-cytotoxic therapies (**Figure 4b-c**), suggesting that cytotoxic chemotherapy provides stronger population bottleneck in HSPCs.

**Figure 4.**
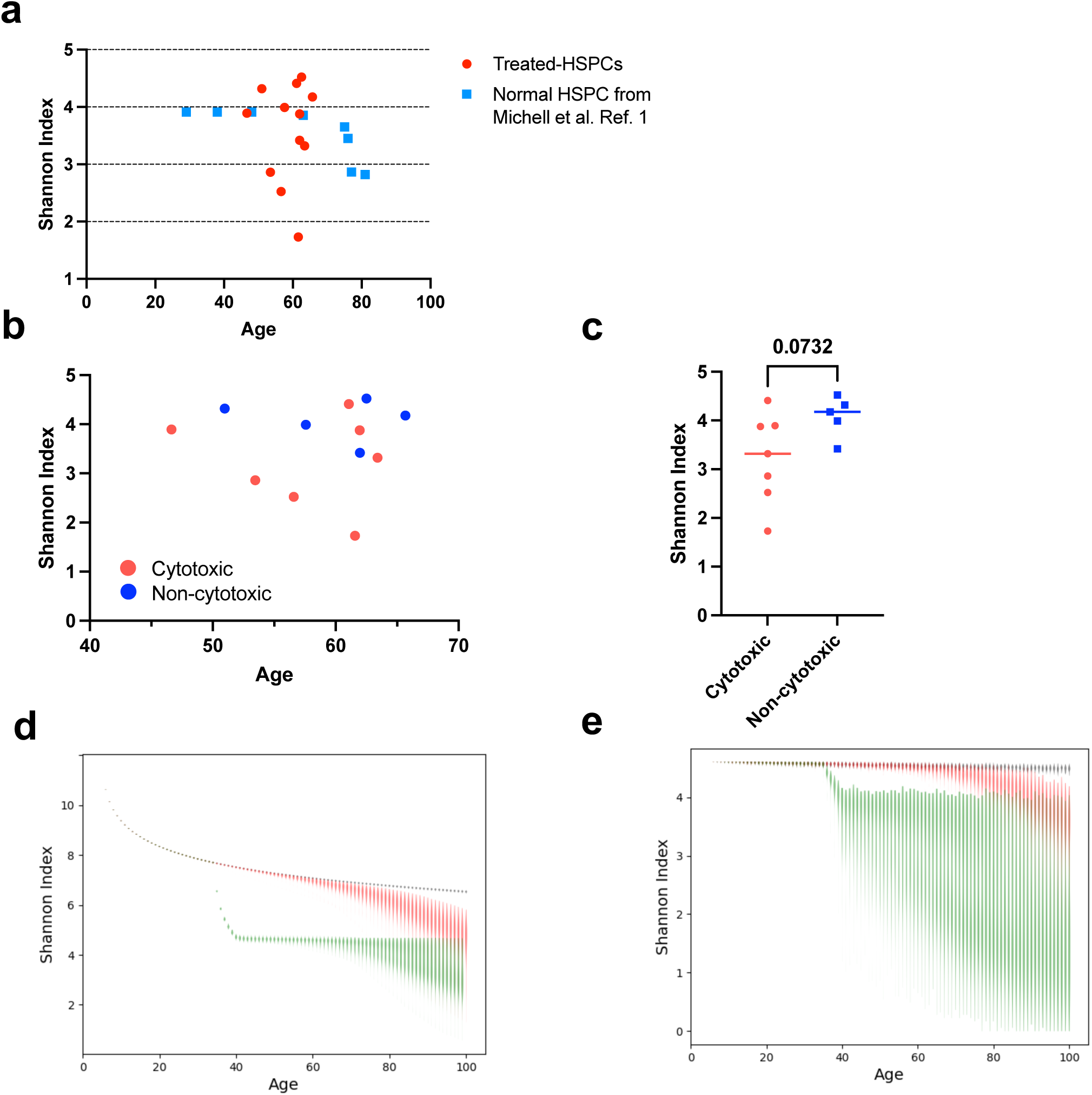
Assessment of clonal diversity in post-chemotherapy HSPCs. (**a**) The Shannon diversity index of post-treatment HSPCs from our cohort compared to indices from normal individuals as reported in Mitchell et al., 2022^1^. (**b**) Comparison of the Shannon diversity index among post-treatment HSPCs from patients who received cytotoxic chemotherapy versus those treated with non-cytotoxic chemotherapy. (**c**) A statistical analysis of the Shannon diversity indices between the two patient groups. P value is obtained from a Mann-Whitney U test. (**d**) The simulation illustrates the projected Shannon diversity index over time for a population of 100,000 HSPCs, modeled with the Moran Model. Each violin plot at each time point represents the results of 100 independent simulations of the model. The black line represents scenarios where acquired mutations do not affect fitness; the red line includes some mutations conferring a selective advantage; the green line indicates the introduction of chemotherapy at approximately ages 35 to 40. (**e**) The simulation displays the Shannon diversity index over time, based on a random sampling of 100 HSPCs from a pool of 100,000, considering the emergence of chemotherapy-resistant mutations (e.g., *TP53* and *PPM1D*).

To better understand the impact of chemotherapy on clonal diversity of HSPCs, we conducted stochastic simulation of HSPC dynamics based on the Moran model^26^. These simulations examined three scenarios: one where all acquired mutations were neutral, another with a mix of neutral and selectively advantageous driver mutations, and a third including the effects of cytotoxic chemotherapy (**Figure 4d).** Consistent with the prior study^1^, in the absence of chemotherapy, the natural occurrence of driver mutations contributed to a decline in diversity index around the seventh decade of life (**Figure 4d**). The introduction of cytotoxic chemotherapy resulted in an immediate and sustained loss in clonal diversity (**Figure 4d**). This pattern was also evident when examining the impact of sample size on the diversity index, particularly when 100 cells were randomly selected from the HSPC pool, which aligns with our experimental conditions (**Extended Figure 5**). Furthermore, in scenarios where chemotherapy-resistant mutations emerge, such as *TP53* and *PPM1D* mutations, the reduction in clonal diversity was exacerbated, with a broader range of variation observed (**Figure 4e**). This is consistent with our experimental observations (**Figure 4a**). Collectively, these findings indicate that cytotoxic chemotherapy accelerates the loss of clonal diversity in HSPCs and its impact persists well beyond the treatment period.

### Clone-specific analysis of driver mutation, mutation rate, and telomere length

The majority of the expanded clades in the post-treatment HSPCs carried driver mutations (30 of 57 clades [53%]), which is in contrast to the previous findings in normal individuals where driver mutations only accompanied 17% (10 out of 57) of the expanded clades^1^ (**Extended Figure 6**). Convergent evolution of DDR pathway mutations (*TP53* and *PPM1D* mutations) was pervasive among treated-HSPCs. For instance, in PID0004, we detected 6 different clades evolving in parallel, each carrying different *PPM1D* mutations (**Figure 3a**). This patient had a second timepoint sample taken 3 years after the first ASCT, which continued to show the convergent evolution of the same *PPM1D* mutations, indicating that all of these *PPM1D*-mutated clones engrafted after ASCT, re-expanded, and re-mobilized (**Extended Figure 7**).

Similarly, in PID0003, PID0005, and PID0008, we observed convergent evolution of multiple different *TP53* mutated clones (**Figure 3a**). These results are indicative of a strong selective pressure from chemotherapy in clonal selection of clones with DDR pathway genes.

In the context of myeloid malignancies, *TP53* mutations frequently co-occur with complex chromosomal aberrations. However, in the post-treatment HSPCs examined in this study, 98% of *TP53*-mutated colonies (106 of 108 colonies) showed normal copy number profiles (**Extended Figure 8**). Only 2 colonies exhibited concurrent chromosomal alterations, specifically a loss of heterozygosity (LOH) on chromosome 17p (PID0002, **Figure 3a, Extended Figure 8**), leading to biallelic alterations in *TP53*. These data suggest that *TP53*-mutated cells do not yet display genomic instability at the CH phase and acquisition of chromosomal aberrations emerge as late-stage leukemogenic events.

To further assess the influence of driver mutations—particularly those in DDR pathway genes— on genomic instability, we compared the mutation rate in wild-type (WT) colonies and clades with or without driver mutations (**Figure 5a and 5c, Extended Figure 9**). We found that some of the clades with *TP53* and *PPM1D* mutations exhibiting significantly higher mutation rate compared to WT colonies (**Figure 5c**). This phenomenon, however, was not universally observed across all clades with driver mutations. Additionally, some of the clades without obvious driver mutations had increased mutation rate. We postulated that the elevated mutation rate in these clades could be attributed to accelerated cellular proliferation rather than the direct consequences of the driver mutations themselves. In corroboration with this hypothesis, some of these clades with high mutation rate demonstrated reduced telomere lengths (**Figure 5b and 5d**). In addition, we observed inverse correlation between mutation rate and telomere length in HSPC colonies further supporting the hypothesis that the increased mutation rate in some of the mutated cells is more likely attributed to accelerated clonal expansion rather than a direct result of the driver mutations per se (**Figure 5e**).

**Figure 5.**
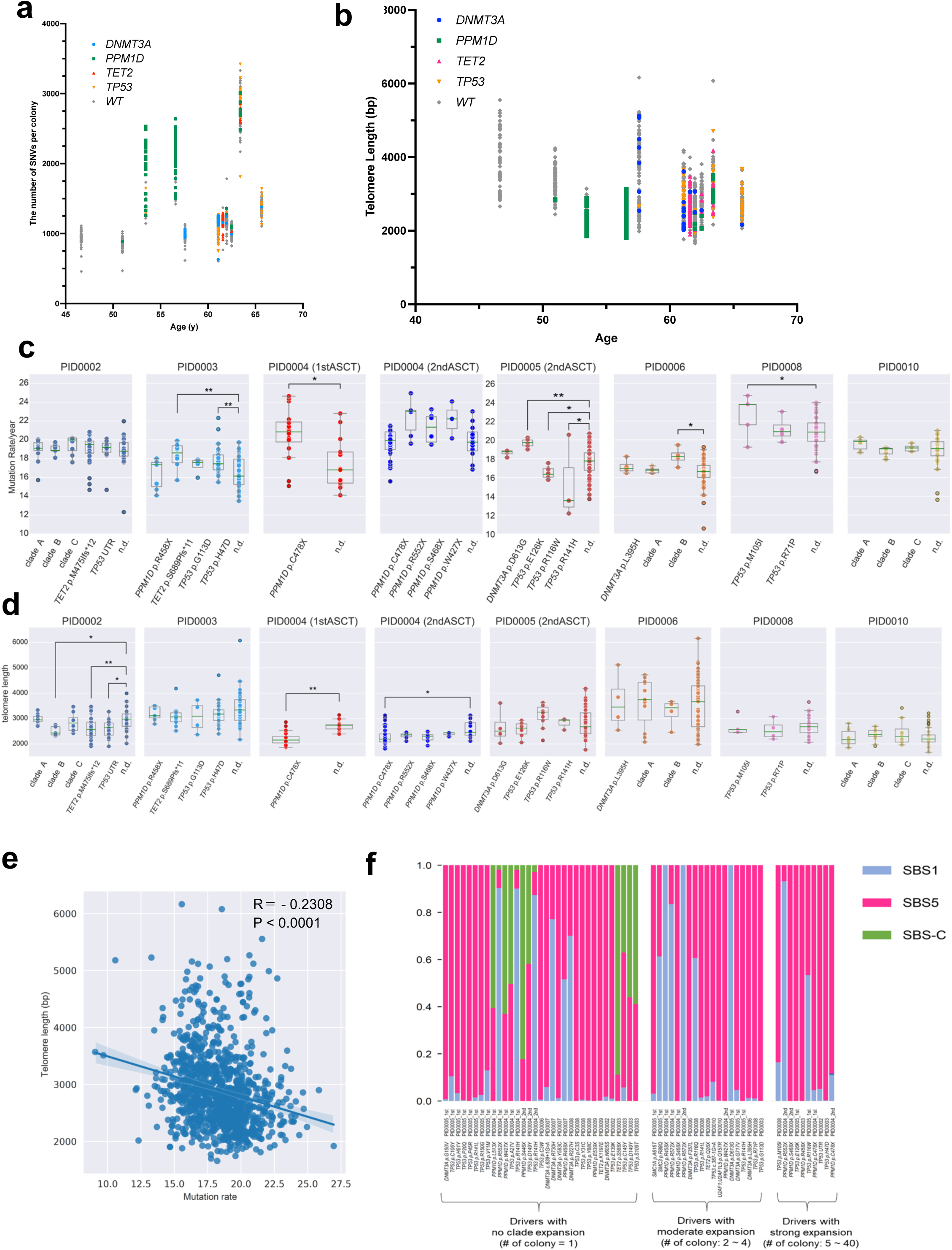
Clone-specific analysis of mutational rate and telomere length. (**a**) Distribution of single nucleotide variants (SNVs) across individual colonies plotted against age, with colonies categorized according to the presence of driver mutations. (**b**) Distribution of telomere length across individual colonies plotted against age, with colonies categorized according to the presence of driver mutations. (**c**) Comparison of mutation rate (SBS1+SBS5 counts per year) between clades with or without driver mutations and wild type (WT) colonies. Statistical significance was assessed using unpaired T-test. * indicates false discovery rate (FDR) < 0.05 and ** indicates FDR < 0.01. The definition of clades without driver mutations is described in **Extended Figure 9**. Clades with fewer than 3 colonies are not shown. n.d.= colonies with no driver mutations. (**d**) Comparison of telomere length between clades with or without driver mutations and WT colonies. Statistical significance was assessed using unpaired T-test. * indicates FDR < 0.05 and ** indicates FDR < 0.01. The definition of clades without driver mutations is described in **Extended Figure 9**. Clades with fewer than 3 colonies are not shown. n.d.= colonies with no driver mutations. (**e**) Scatter plot correlating mutation rate (SBS1+SBS5 counts per year) and telomere length in each colony. (**f**) Assessment of the contribution of specific mutation signatures on individual driver mutations detected in treated colonies. Mutations are segregated based on the clade expansion. All SBS-C related mutations were found in colonies with no clade expansion.

Next, given the evidence of chemotherapy-induced somatic mutations in HSPCs by chemotherapy, particularly by melphalan, we investigated the potential contribution of chemotherapy in directly causing any of these driver mutations. To estimate the contribution of specific mutation signatures to the development of driver mutations, we mapped the nucleotide context of these mutations onto each signature following established methods (**Figure 5f**)^27^.

Predominantly, the driver mutations corresponded with signatures SBS-1 or SBS-5. However, the *TET2* p.S588X mutation in PID0003 and the *PPM1D* p.S446X mutation in PID0004, identified at the second timepoint, had an 89% and 82% probability of contribution from SBS-C, respectively. Furthermore, additional driver mutations—including two *TP53* mutations in PID0003 and one *PPM1D* plus one *TP53* mutation in PID0004, detected at the first timepoint— exhibited over a 50% likelihood of association with SBS-C. Notably, the mutations linked to SBS-C did not demonstrate clonal expansion in our phylogenetic analysis (**Figure 3 and Figure 5f**). This implies that while melphalan treatment may induce some driver mutations in HSPCs, these mutations do not appear to confer a substantial selective advantage, potentially due to their induction later in life which limits the opportunity for clonal expansion.

### Mapping the clonal origin of therapy-related myeloid neoplasms in PBSC samples

In the current cohort, nine out of ten patients developed t-MNs with a median of three years (range: 1-8 years) following peripheral blood stem cell (PBSC) collection. We obtained bulk DNA from bone marrow samples at t-MN diagnosis and conducted bulk WGS (median 52x coverage). The mutational profiles and chromosomal abnormalities found in t-MN samples are detailed in **Table 2**.

**Table 2.** The list of driver mutations and copy number alterations (CNAs) detected in therapy-related myeloid neoplasms (t-MN) samples by bulk whole genome sequencing.

| Sample ID | Driver 1 | VAF | Shared with colonies* | Driver 2 | VAF | Shared with colonies* | CNAs |
| --- | --- | --- | --- | --- | --- | --- | --- |
| PID0001 | *GNAS*<br>p.R186C | 0.25 | No | - | - | - | Amp 3q, Del 11q |
| PID0002 | *TP53*<br>UTR5<br>mutation | 0.885 | Yes | *TET2*<br>p.M475Ifs*12 | 0.5 | Yes | Del 5q, -7, Del 17p |
| PID0004 | *TP53*<br>p.C3W | 0.92 | No | - | - | - | Del 5q, Del 6q, 17p CN-LOH |
| PID0005 | *TP53*<br>p.E126K | 0.45 | Yes | *DNMT3A*<br>p.Y722X | 0.26 | Yes | Del 5q, -7, Del 17p, -18 |
| PID0006 | *TP53*<br>p.C3W | 0.833 | Yes | - | - | - | Poor sample quality |
| PID0007 | *TP53*<br>p.V173M | 0.767 | No | - | - | - | Del 5q, Del 17p |
| PID0008 | *TP53*<br>p.M105I | 0.652 | Yes | *TP53* splice site<br>mutation | 0.25 | Yes | Del 7q, Del 17p, -21 |
| PID0009 | *TP53*<br>p.F95C | 0.436 | No | *TP53*<br>p.G22Afs*16 | 0.293 | No | -3, -4, Del 19p, -21 |
| PID0010 | *WT1*<br>p.R368Tfs<br>*5 | 0.448 | No | - | - | - | Normal |

\*Mutations shared with corresponding colonies are also indicated.
**Abbreviations:** VAF: variant allele frequency, CNAs: copy number alterations, CN-LOH: copy neutral loss of heterozygosity

To understand the clonal origin and evolutionary history of t-MN development, we performed phylogenetic analysis integrating the genomes of HSPC colonies and t-MNs. This analysis identified most recent common ancestor (MRCA) of t-MNs in 5 of 9 (56%) patients’ PBSC samples through the analysis of shared variants between individual colonies and t-MN genomes (PID0002, PID0005, PID0006, PID0008, and PID0010, **Figure 6, Extended Figure 10**).

**Figure 6.**
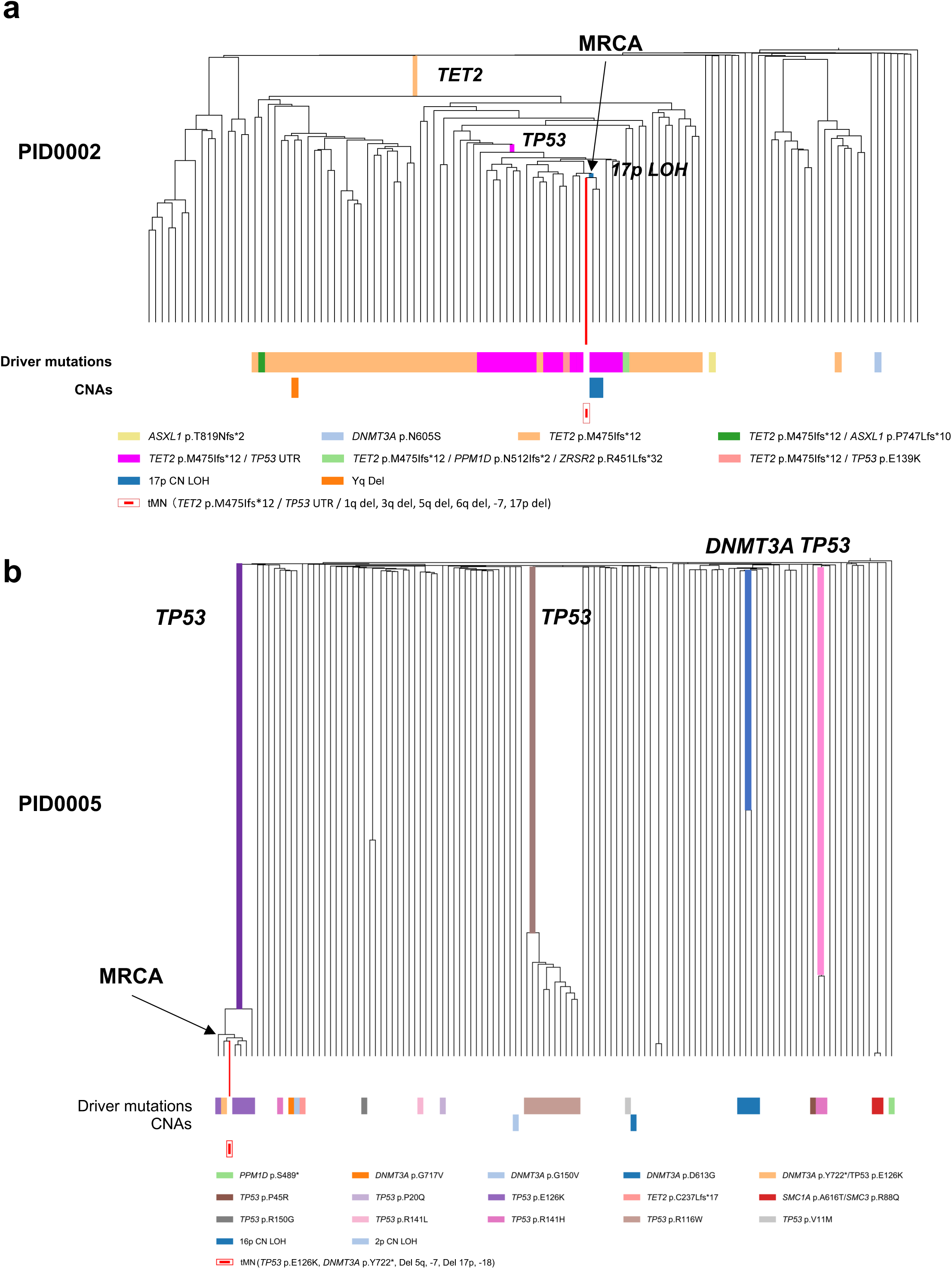
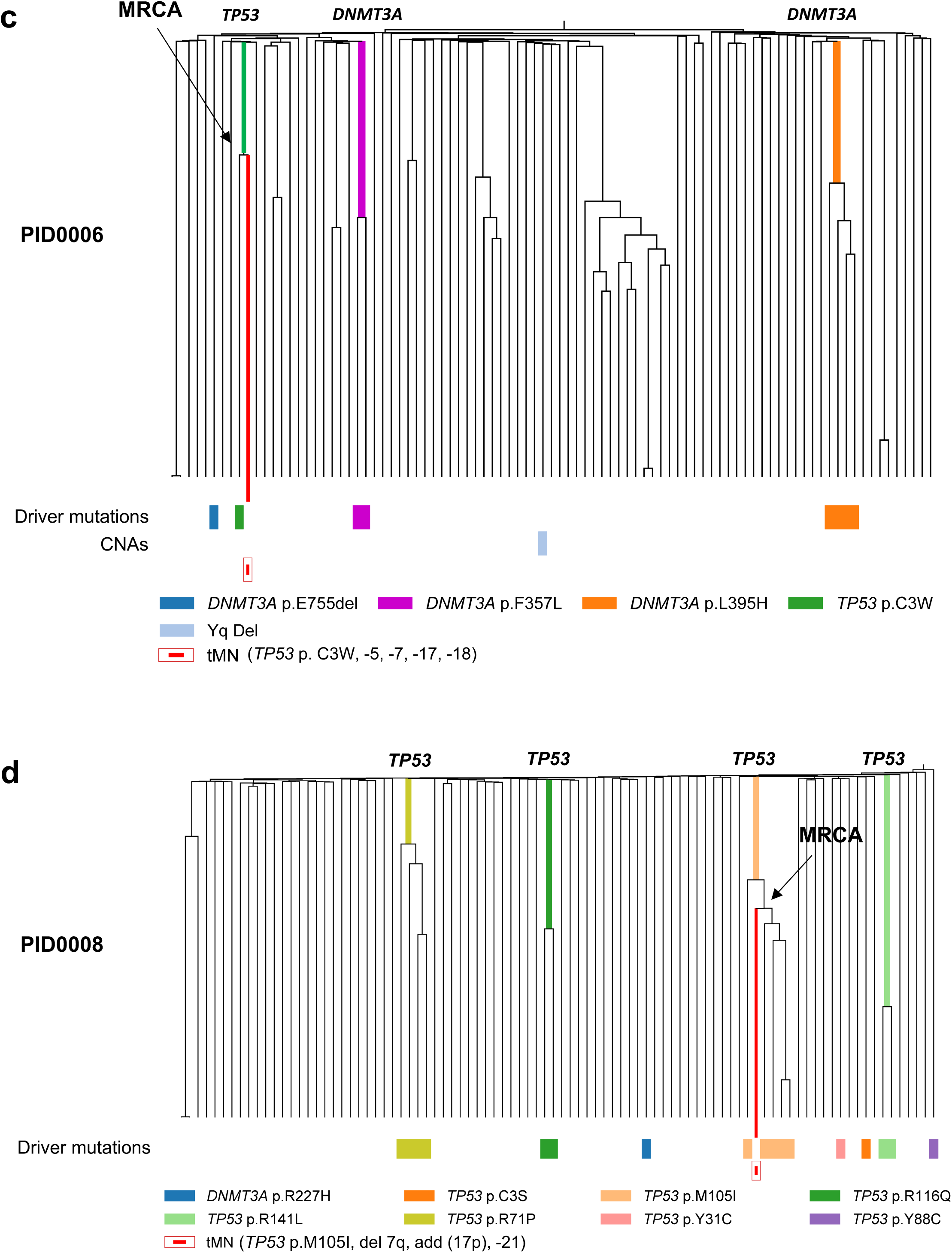
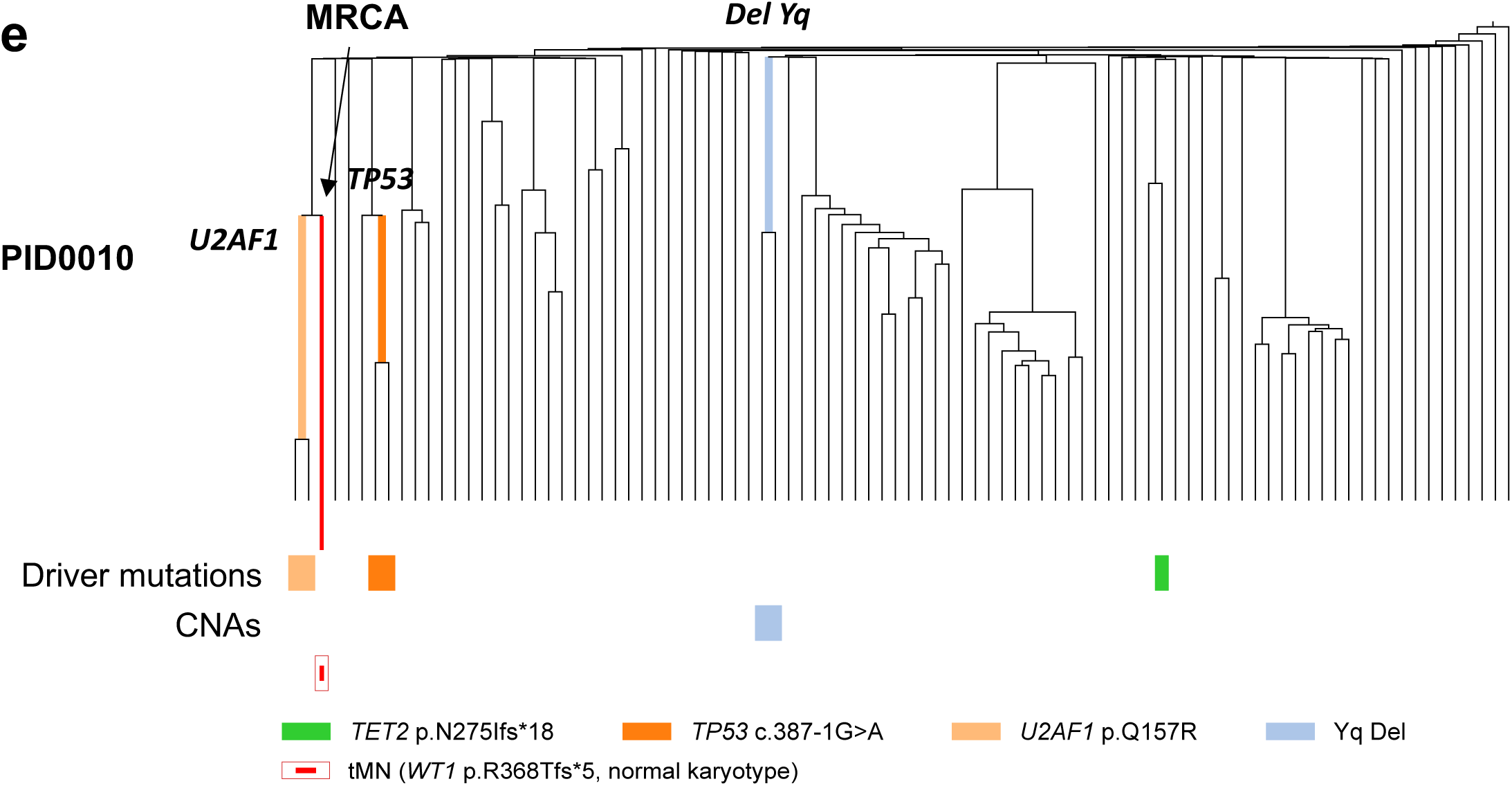
Phylogenetic relationships between post-treatment HSPCs and corresponding t-MN samples. (**a**) Integrated phylogenetic tree for PID0002 highlighting the most recent clonal ancestor (MRCA) pinpointed to a clone with concurrent *TET2*, *TP53* mutations, and 17p loss of heterozygosity (LOH). (**b**) Integrated phylogenetic tree for PID0005 where MRCA was identified in *TP53* and *DNMT3A* mutated clone. (**c**) Integrated phylogenetic tree for PID0006 where the MRCA is identified within a clone possessing a *TP53* mutation. (**d**) Integrated phylogenetic tree for PID0008 with the MRCA traced to a *TP53*-mutated clone. (**e**) Integrated phylogenetic tree for PID0010 showing the MRCA located at a branching point preceding the acquisition of a *U2AF1* mutation.

PID0002 developed therapy-related myelodysplastic syndrome (t-MDS) one year post-ASCT. Analysis of this patient’s HSPC colonies revealed a prominent clone with a *TET2* mutation, containing subclones with an additional *TP53* mutation. Within these subclones, two colonies also possessed 17p loss of heterozygosity (LOH), indicative of biallelic *TP53* alterations. The MRCA of the t-MDS clone was traced to the branchpoint of these specific clones, which carried mutations in *TET2*, *TP53*, and 17p LOH (**Figure 6a, Extended Figure 10a**). The t-MDS sample also presented with additional copy number alterations (CNAs) such as monosomy 5 and 7, that were absent in the HSPC colonies, indicating subsequent acquisition post-MRCA.

PID0005 underwent two PBSC collection 20 and 6 years prior to the development of t-MDS, respectively. PBSC colonies at the first timepoint showed polyclonal HSPCs without any CH mutations. However, the second timepoint samples exhibited multiple parallel clones with *TP53* mutations and *DNMT3A* mutations. Notably, one clone with the *TP53* p.E126K mutation underwent significant expansion and included single colony with an additional *DNMT3A* p.Y722* mutation. We traced MRCA of the t-MDS clone adjacent to the colony with this double *TP53* and *DNMT3A* mutations (**Figure 6b, Extended Figure 10b**). Bulk sequencing of the t-MDS sample identified both mutations with additional CNAs involving chromosome –5, –7, –17p, and –18.

PID0006’s t-MDS emerged eight years after PBSC collection. This patient’s HSPC colonies comprised three distinct clones with different *DNMT3A* mutations and one colony with a *TP53* p.C135W mutation without any CNAs. The t-MDS sample harbored the same *TP53* mutation but also monosomy 17, indicating biallelic *TP53* alteration. Phylogenetic integration pinpointed the MRCA of the t-MDS clone with the colony harboring the *TP53* mutation (**Figure 6c, Extended Figure 10c**).

PID0008 presented with t-MDS six years post-collection. Multiple parallel *TP53*-mutated clones were identified in HSPC colonies without any concurrent CNAs. Among them, the *TP53* p.M237I mutation was also found in the t-MDS sample. The MRCA was located within the clones carrying this *TP53* mutation (**Figure 6d, Extended Figure 10d**). Additionally, the t-MDS sample had an essential splice site *TP53* mutation not detected in the colonies, indicating that the second *TP53* mutation was acquired post-MRCA. The shared feature of these 4 cases is that sequential acquisition of secondary mutations and biallelic alteration of *TP53* is the critical factor determining malignant transformation.

The 5^th^ case, PID0010, developed t-MDS three years after PBSC collection. The patient’s HSPC colonies included two with *U2AF1* mutation and two with *TP53* mutations, none of which appeared in the t-MDS sample. However, phylogenetic analysis traced the MRCA of t-MDS to a branchpoint preceding the *U2AF1* mutation, indicating the clone that led to t-MDS had diverged prior to acquiring *U2AF1* (**Figure 6e, Extended Figure 10e**). Instead, the t-MDS sample contained a *WT1* mutation.

In other 4 cases (PID0001, PID0004, PID0007, and PID0009), phylogenetic analysis did not identify an MRCA in the studied colonies (**Extended Figure 11 and 12**). The t-MN clones in these patients diverged early and shared few mutations with HSPC colonies. This was unexpected, as some colonies exhibited high-risk CH mutations such as *TP53* and *PPM1D*. Given the limited number of analyzed colonies (∼100 per patient), these results may reflect sampling bias. Alternatively, t-MNs may have originated from residual bone marrow HSPCs not mobilized to PBSCs. To explore this hypothesis, we examined the mutation signatures in the t-MN samples, positing that if t-MNs arose from these residual HSPCs, they would carry the mutational footprint of exposure to the high-dose melphalan used in the conditioning regimen. Corroborating this theory, approximately 40% of the mutations in PID0007’s and PID0009’s t-MN sample bore the SBS-C signature, which is indicative of melphalan exposure (**Extended Figure 13**). Given that PID0007 and PID0009 did not receive melphalan during the induction phase, it suggests that the t-MN originated from bone marrow HSPCs that were subjected to high-dose melphalan during the conditioning regimen. While these findings require validation in a larger cohort, they suggest two distinct developmental pathways for t-MNs post-ASCT: while most arise from clones within PBSCs, a minority may develop from non-mobilized bone marrow HSPCs that underwent conditioning therapy (**Extended Figure 14**).

## Discussion

The clinical toxicity of cancer chemotherapy can manifest in the hematopoietic system, ranging from transient myelosuppression to the devastating development of t-MNs. Here, we utilized a single-colony WGS approach to evaluate the influence of cancer chemotherapy on the genome and clonal dynamics of human HSPCs. While the list of chemotherapies evaluated in this study is by no means exhaustive, we found that melphalan treatment resulted in the induction of treatment-related mutations in HSPC genomes, including several driver mutations. Despite sharing its classification as an alkylating agent with melphalan, cyclophosphamide-treated HSPCs did not manifest treatment-induced mutations. This discrepancy may be explained by the ability of HSCs to metabolize cyclophosphamide into an inactive formulation via the activity of ALDH^24^. This aligns with the well-established use of cyclophosphamide in clinical practice, where it is safely administered to prevent graft-versus host disease three days after the infusion of allogeneic HSCs without detrimentally affecting engraftment (also known as post-Cy regimen)^28^. Furthermore, this is consistent with a significantly reduced incidence of t-MNs in breast cancer patients treated with cyclophosphamide, as compared to those treated with melphalan^29^.

Beyond its mutagenic implications, cancer chemotherapy appears to have a strong influence on the clonal architecture and dynamics of HSPCs. In comparison with HSPCs from healthy untreated individuals that were published previously^1^, treated-HSPCs exhibited marked loss of clonal diversity, which was driven by the expansion of multiple clones harboring driver mutations such as *TP53* and *PPM1D* mutations. The clonal heterogeneity of chemotherapy-treated HSPCs from patients in their 40s and 50s mirrored that typically observed in healthy individuals in their 70s or beyond, suggesting that chemotherapy might accelerate the genetic ageing of HSCs. Thus, the temporal repercussions of chemotherapy extend beyond immediate cytotoxic effects, insinuating a long-term reshaping of the clonal architecture of HSC pools that parallels the natural genetic evolution associated with ageing.

One such long-term consequence of chemotherapy treatment is the development of t-MNs. In our study, through comparative genomic analysis of t-MNs and individual HSPCs, we traced the clonal origin of t-MNs to a specific HSPC clone. While prior investigations from our team and others have demonstrated that t-MN driver mutations are detectable in the blood drawn years ahead of t-MN development^30,31^, the single-cell analysis incorporated in this study revealed a more complicated picture of clonal evolution from normal HSPCs to t-MNs. We observed that chemotherapy-treated HSPCs frequently undergo branching convergent evolution, generating multiple parallel clones with *TP53* mutations, each possessing high-risk leukemic potential in principle (**Figure 7**)^32,33^. Intriguingly, among the multitude of parallel clones with *TP53* mutations, only one typically progresses to a therapy-related myeloid neoplasm (t-MN), raising a question regarding the underlying genetic or epigenetic determinants for this clonal selection. A noteworthy observation from our analysis is that clones acquiring biallelic *TP53* alteration (with a second mutation or through LOH) and additional chromosomal alterations seemingly possess the highest leukemic potential, which is consistent with the known high prevalence and poor prognosis of biallelic *TP53* alterations in t-MNs^20,34^. These findings underscore the critical role of secondary mutations in dictating clonal selection and malignant transformation in t-MNs (**Figure 7**). Clinically, this observation highlights the importance of monitoring not only the clonal proliferation of *TP53* mutation by bulk sequencing but also the careful evaluation of allelic status, which likely requires single-cell genotyping analysis. This approach will be instrumental in the precise early identification of the clones poised to evolve into to t-MNs.

**Figure 7.**
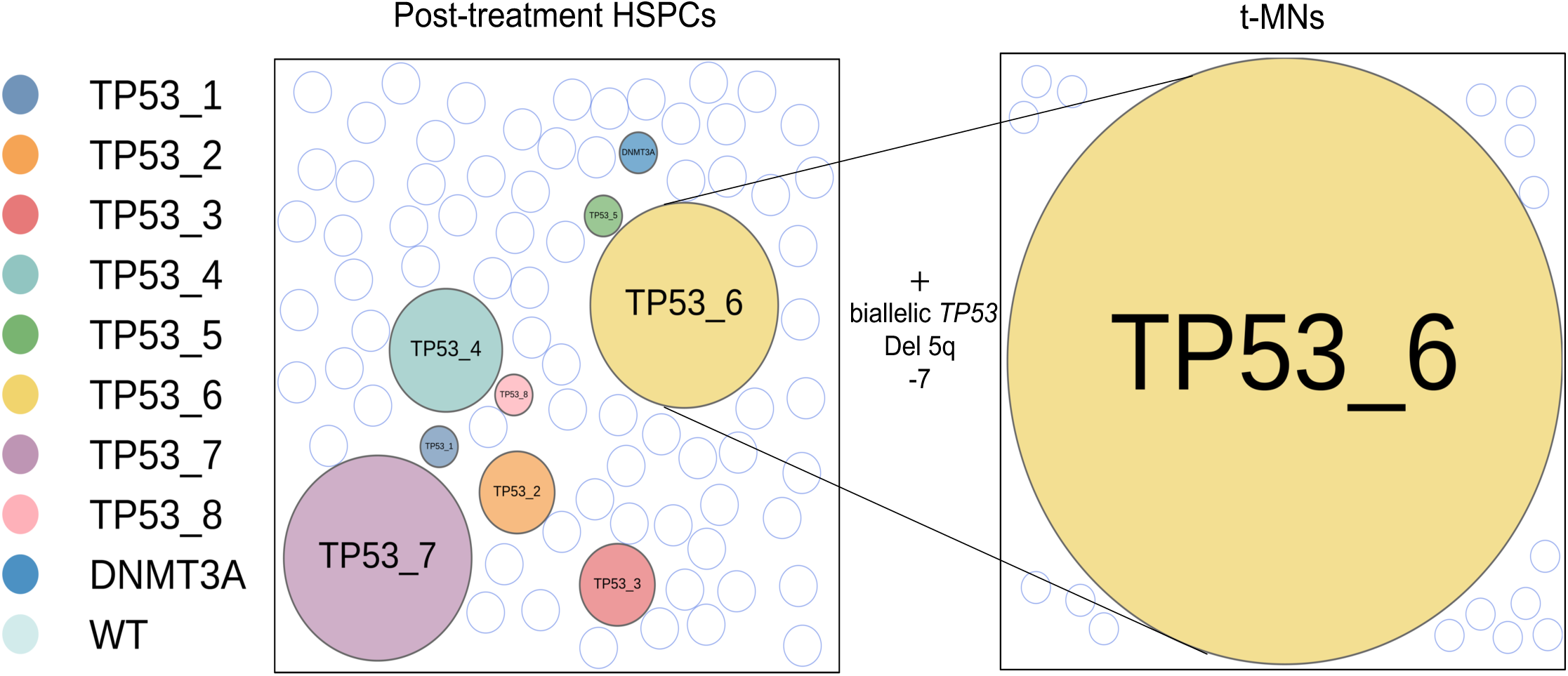
Clonal evolution of t-MNs. Illustrative representation of clonal evolution from post-chemotherapy HSPCs to t-MNs (PID0008). The figure portrays the parallel evolution of multiple HSPC clones, each harboring different *TP53* and *DNMT3A* driver mutations of varying clonal sizes, interspersed with wild-type (WT) cells. It highlights one clone that develops biallelic *TP53* alterations and acquires additional copy number alterations, ultimately undergoing selective clonal expansion with the transformation to t-MNs.

In the majority of cases, our integrated phylogenetic analysis traced the clonal origins of t-MNs to PBSCs. However, in a subset of cases, we were unable to determine the clonal origins within PBSCs. Notably, in two of those case, the t-MN samples displayed a mutational signature indicative of exposure to conditioning melphalan chemotherapy, pointing to an origin from bone marrow HSPCs that were not part of the mobilized PBSC population. These results are consistent with the findings by Diamond et al. that subset of post-ASCT t-MNs exhibit melphalan signatures^35^. Based on these findings, we propose the existence of two distinct developmental pathways for t-MNs following ASCT: one originating from transplanted PBSCs and another from non-mobilized bone marrow HSPCs. Clinically this implies that screening PBSC samples for high-risk CH mutations may not always predict the emergence of t-MNs.

In summary, our study revealed the differential effects of cancer chemotherapy on the genome of HSPCs, underscoring the need for future research involving a more systematic evaluation of chemotherapy’s impact on the HSC genome and its association with long-term consequences. We also found that chemotherapy accelerates the loss of clonal diversity in HSPCs by promoting the expansion of multiple clones harboring DDR pathway mutations, notably *TP53* and *PPM1D*. Among these, one clone eventually transforms into t-MNs. Further investigation is warranted to identify the precise genetic or epigenetic determinants dictating this transformation and clonal competition. Additionally, establishing clinical strategies to monitor patients exposed to high-risk chemotherapy is crucial for the early detection and potential interception of emergent t-MNs.

## Methods

### Patient samples

Aliquots of cryopreserved peripheral blood stem cells (PBSCs) from multiple myeloma patients who underwent autologous stem cell transplantation (ASCT) were utilized to culture single HSPC colonies. For those patients who subsequently developed t-MNs, genomic DNA was extracted from their bone marrow mononuclear cells for WGS analysis. Written informed consent for the collection and analysis of samples was obtained from all participating patients. The study protocols were conducted in accordance with ethical guidelines and received approval from The University of Texas MD Anderson Cancer Center’s institutional review board.

### Single-HSPC colony formation and DNA extraction

Cryopreserved PBSCs were carefully thawed and then cultured at 37°C using MethoCult™ H4435 Enriched Medium (STEMCELL Technologies Inc.), adhering to the manufacturer’s instructions. After a period of 14-18 days of incubation, colonies were identified using microscope, and individual single-cell-derived colonies were harvested and suspended in phosphate-buffered saline (PBS). Genomic DNA was then extracted from these colonies using the DNeasy Blood & Tissue Kit (Qiagen, 69506), following the protocol provided by the manufacturer. The quality of the extracted DNA was assessed using Qubit fluorometric quantitation and/or TapeStation analysis prior to WGS.

### Whole genome sequencing (WGS) and read alignment

WGS was performed in the Advanced Technology Genomics Core (ATGC) Facility at MD Anderson Cancer Center. Briefly, Illumina compatible, uniquely dual-indexed libraries were prepared from 200ng of RNase treated DNA. The DNA was sheared to approximately 350bp using Diagenode Biorupter Pico, then libraries were prepared using the KAPA Hyper Library Preparation Kit (Roche Sequencing Solutions, Inc.). The libraries were amplified with 3-8 cycles of PCR, then assessed for size distribution using the 4200 TapeStation High Sensitivity D1000 ScreenTape (Agilent Technologies) and quantified using the Qubit dsDNA HS Assay Kit (Thermo Fisher). Equimolar quantities of the indexed libraries were multiplexed, 31-32 libraries per pool. The pool was then quantified by qPCR using the KAPA Library Quantification Kit (KAPA Biosystems) and sequenced on the Illumina NovaSeq6000 S4-300 flow cell using the 150nt paired end format.WGS with low input material was carried out using Illumina NovaSeq with 150 bp paired-end sequencing, which provided approximately 28x coverage per colony sample. Quality control for the raw sequence data, including trimming of adapters, was conducted using Trim Galore, version 0.6.5. The sequencing reads were then aligned to the human reference genome (NCBI build 37) using the BWA-MEM algorithm. Sequence deduplication was carried out employing the Picard MarkDuplicates function.

### Somatic mutation calling

Variant calling was conducted using the Genome Analysis Toolkit (GATK) Mutect2 (ver 4.2.0.0) against a pool of unmatched normal control samples^36^. Post Mutect2 filtration, additional custom filters were applied to refine mutation calls, aiming to eliminate germline variants, sequencing errors, artifacts, misalignments, and mutations acquired during the culture. To filter germline variants, we adopted the methodology used by Yoshida et al.^10^. Briefly, we fitted a binomial distribution model to Mutect2-filtered variants and total depth across all samples from a single patient. A one-sided exact binomial test was performed with the null hypotheses that the variants would display a binomial distribution with a probability of 0.5 (0.95 for sex chromosomes in male). Variants with P-value greater than 10e-10 were classified as germline. Additional variant filtering was conducted following the method described by Lee-Six et al.^4^, and consists of; 1) Removal of mutations located within 10 base pairs of each other, 2) Mutations exhibiting sequencing coverage of fewer than five reads for autosomes or for X chromosomes in females and fewer than three reads for sex chromosomes in males were assigned a status of ‘NA’ in the respective sample. To minimize the risk of sequencing artifacts, any mutations bearing this ‘NA’ status identified in more than five samples within a single patient were excluded from the dataset, 3) Removal of mutations with a mean VAF across all samples containing mutant reads of 0.3 or less, and 4) Removal of variants with a VAF below 0.1 observed in over 10% of samples.

### Copy number analysis

Copy number profiles were generated from WGS data using AscatNGS pipeline (https://github.com/cancerit/ascatNgs). One of the colonies that belonged to the unexpanded clades from the same patients was used as a matched normal. These called regions that have larger than 5Mbp in this process are considered true copy number aberrations (CNAs).

### Estimation of telomere length from WGS data

Based on the WGS data from each colony, telomere length was estimated using Telomerecat algorithm^37^.

### Mutation signature analysis

We applied SigProfiler to our WGS data^38^ and cross referenced the extracted signatures with the known COSMIC signatures (https://cancer.sanger.ac.uk/cosmic/signatures/SBS/). Mutation signatures that show cosine similarity of 0.80 or lower were considered as novel signatures. Subsequently, both novel and previously identified signatures were reassigned to the samples, with careful consideration of each patient’s clinical history. To confirm the presence of each mutated signature in the samples and estimate its contribution, fitting algorithms from SigProfilerExtractor^39^ and mmSig^40^ were utilized, which showed consistent results.

### Phylogenetic tree reconstruction

Somatic mutations on all samples were described in the PHYLIP format files, and the alleles on the genomic positions where the variants were not called were considered to be reference alleles. The putative allele sequences of the hypothetical fertilized zygote, which consist of reference alleles after removal of the germline mutations, were also added as an outgroup of the samples. The phylogenetic trees for these samples are inferred as follows.

PHYML (ver 3.3.20211231) was used to infer the cladogram based on the maximum-likelihood method. The number of the bootstrap to estimate the initial cladogram for the samples was 100. The node between the outgroup and other samples is set as root. After the copy number variations analysis, PHYLIP files for each clade on the initial phylogenetic tree were created, and the alleles that existed on the loss-of-heterozygosity (LOH) region of more than a single sample were removed. Then, the cladogram for the clade was re-evaluated with PHYML with the bootstrap number as 1000. These reevaluations of cladogram were performed for each clade in descending order.

### Branch length adjustment on phylogenetic tree

Based on the cladogram, ancestral sequence reconstruction (ASR) was performed for all ancestry nodes with the phangorn library in the R language. Mutational signature analysis was conducted on the somatic mutations identified in colony samples and the inferred ancestral sequences. This analysis aimed to assess the relative impacts of intrinsic and extrinsic factors on each node, quantifying their respective influences. Then, we evaluated the differences in the signature counts between each pair of a child node and a parent node by subtracting the signature counts on the parent node from the child node. If a negative value was computed for any signature count on the branch, two criteria were independently assessed for each branch:

1) Whether the value fell within the range of –1 to 0, or
2) Whether the absolute value of the negative value was less than 5% of the total signature count for the branch.

When either condition was met, the negative value was regarded as an artifact. All the negative values on each branch in our phylogram met either 1st or 2nd criteria, and these negative counts were consequently treated as zero.

After this process, the sum of the counts of SBS1 and SBS5, both clock-like signatures, is applied to the branch length to represent the evolutionary time for these phylograms.

### Simulation of HSPC population dynamics and changes in clonal diversity

We adopted the Moran model to represent a turnover of a healthy HSPC population over age with the accumulation of mutations, where a cell for death is randomly chosen and another cell is randomly chosen for division according to the fitness proportionate selection during one time step^26^. This model keeps the number of the HSPC population constant. Let us begin with *N* active HSPCs after tissue maturation and assume that one time step corresponds to 1/*N* year. Then, each HSPC divides once per year on average. We assumed that the HSPC population is established at 5 years old, as previously described^4^, and each cell is defined as an independent clone in the initial population. As time passes, some clones may expand in the HSPC population and others may go extinct, leading to the reduction of the clonal diversity. Initially all HSPCs have the same fitness, 1.0, but their fitness additively increases with *s* when a driver mutation occurs during a cell division, where the mutation rate per division is denoted by *u*. Therefore, the probability that the *k*-th cell with *i_k_* driver mutations is chosen for division is given by

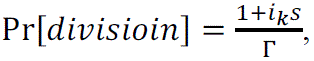

where 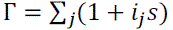. When *s* = 0, the probability of division is the same among all cells irrespective of their mutations, indicating that the population dynamics is neutral. Note that the probability that a cell is chosen for death is the same among all HSPCs in this model.

Next, we introduced the effect of chemotherapy in HSPC population. The chemotherapy damages HSPC population to decrease the total population number into εN (ε < 1) during the chemotherapy. After the treatment, the total number is assumed to be recovered within β years. We consider two different scenarios for the chemotherapy,

1) The probability that a cell is chosen for death is the same among all HSPCs as in the healthy state.
2) Cells that have a specific mutation are resistant to death during the chemotherapy.

In the latter case, the mutation resistant to death by the chemotherapy would emerge before or during the chemotherapy with probability *m* per division. The probability that the *k*-th cell is chosen for death during the chemotherapy is given by 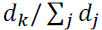, where 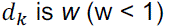 for cells that have the resistant mutation, and 1.0 for cells that don’t have the resistant mutation. Simulation codes are attached in supplemental data.

## Data Availability

Whole genome sequencing data from all colonies are available at Sequence Read Archive (SRA) with the project number PRJNA1058953.

## Acknowledgments

This study was supported by National Institutes of Health Grants R01CA237291 (MG and KT), R01CA262636 (CD, KB, and KT), and P01CA265748 (MG and KT), Physician Scientist Program at MD Anderson (to KT), Andrew Sabin Family Foundation Award (to KT), American Society of Hematology Scholar Award (to KT), Dresner Foundation Early Investigator Award (to KT), Leukemia and Lymphoma Society Scholar Award (KT), Break Through Cancer Foundation (KT and GGM), Lyda Hill Foundation (to PAF), the Charif Souki Cancer Research Fund (to HK), the MD Anderson Cancer Center Leukemia SPORE grant (NIH P50 CA100632) (to HK and KT), the MD Anderson Cancer Center Support Grant (NIH/NCI P30 CA016672), and generous philanthropic contributions to MD Anderson’s Moon Shot Program (to PAF, KT, GGM, and HK). RZO, the Florence Maude Thomas Cancer Research Professor, would like to acknowledge support from the Dr. Miriam and Sheldon G. Adelson Medical Research Foundation.

## Author contributions

KT designed the study, provided leadership, managed the study team, and wrote the manuscript. HU performed experiments and bioinformatics analysis. KS and HH performed simulation study. CDK, KF, LZ, JZ, and XS performed bioinformatics analysis. JN and MC helped data analysis and study conception. JIH, ZC, SC, TT, ZL, YH, ET, HT, LL, and CG performed experiment. CD, ND, NP, FR, RZO, MQ, KB, HK, RKS, CBR, ES, and GGM provided samples and treated patients. SC, DN, PAF, and MG provided critical scientific input and assisted study conception. All authors read and approved the manuscript.

---

**Extended Figure 1.**
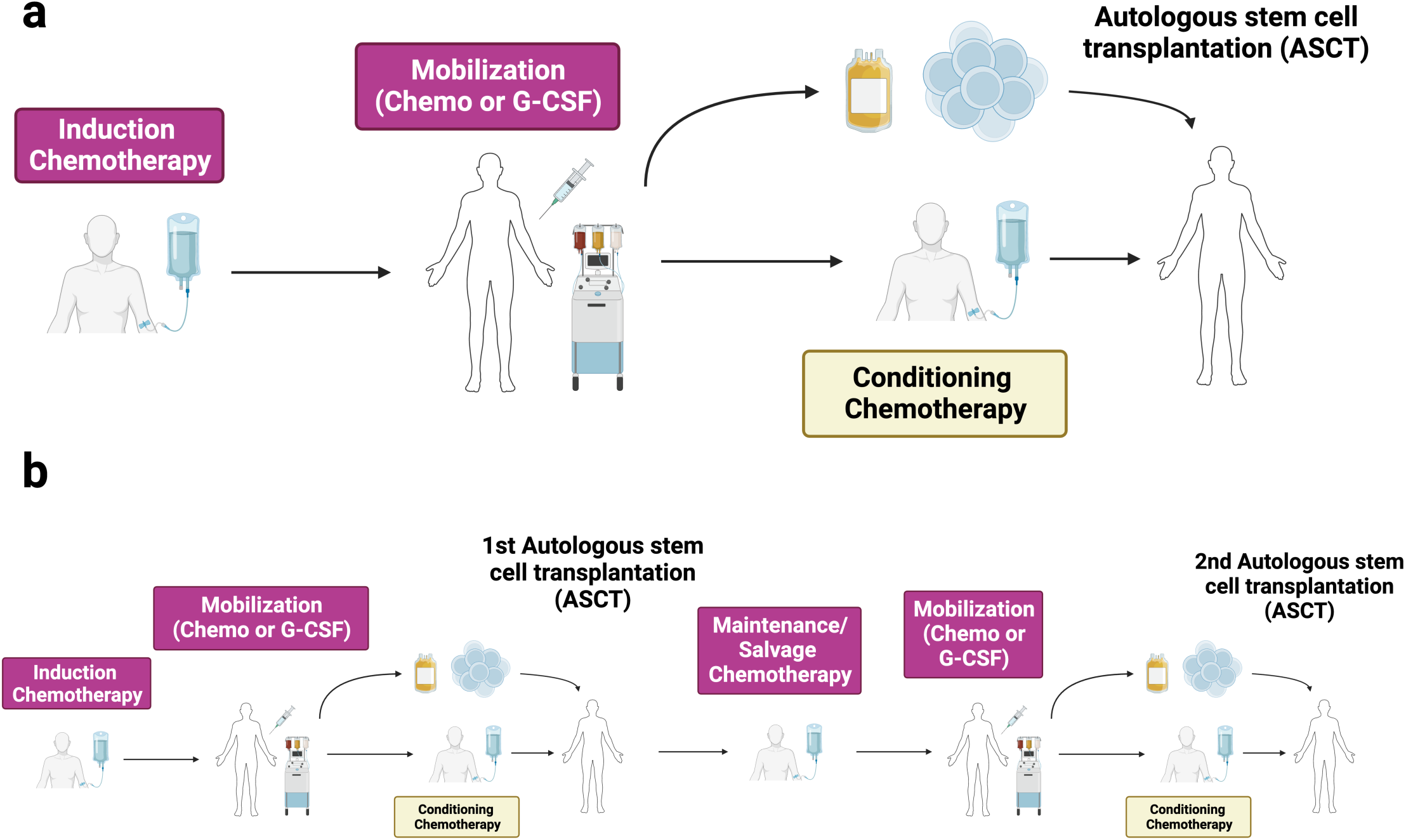
Visual summary of chemotherapy exposure for mobilized peripheral blood stem cells (PBSCs). (**a**) Diagram depicting the chemotherapy exposure of PBSCs from patients undergoing a single autologous stem cell transplant (ASCT), which includes agents administered during induction therapy and any mobilization regimen. (**b**) For patients receiving two ASCTs, this figure outlines the additional exposure of second-timepoint PBSCs to chemotherapies administered during induction, the first mobilization, any maintenance or salvage therapies post-first ASCT, and the mobilization for the second ASCT. Notably, in both scenarios, PBSCs do not encounter chemotherapies applied during the conditioning phase of the transplant.

**Extended Figure 2.**
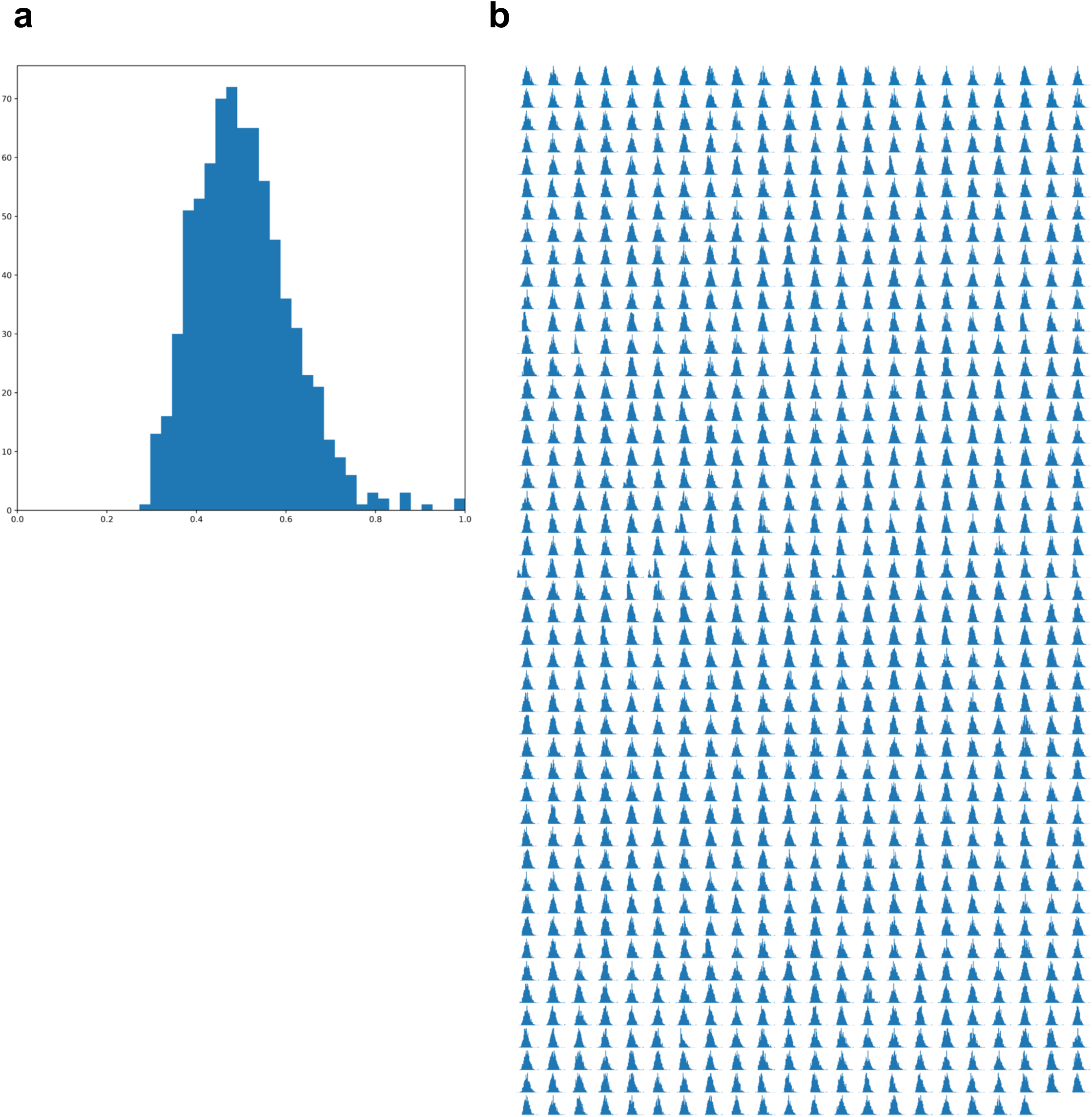
Analysis of variant allele frequency (VAF) for somatic mutations in HSPC colonies. (**a**) A typical VAF histogram for a single colony, displaying a pronounced peak at 50% VAF, which is indicative of its single-cell origin. (**b**) Aggregate VAF distribution across all sequenced colonies, totaling 1,032, illustrating the frequency of somatic single nucleotide variants (SNVs) detected

**Extended Figure 3.**
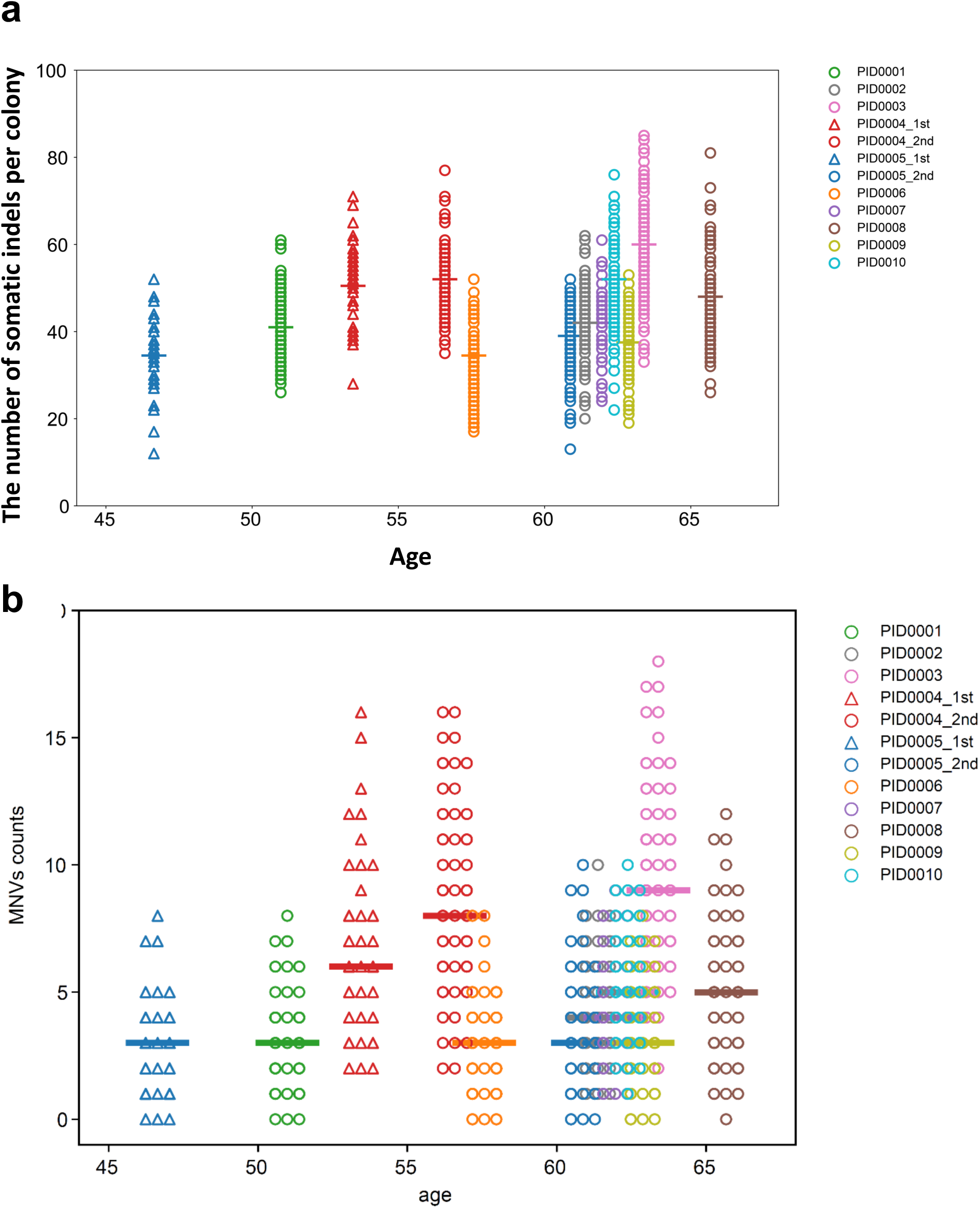
Age-related distribution of somatic indels and multiple nucleotide variants (MNVs) in HSPC colonies. (**a**) Graph showing the correlation between the number of somatic indel variants in each colony and the age of the individual from whom the colony was derived. (**b**) Graph depicting the relationship between the number of somatic MNVs per colony and the age.

**Extended Figure 4.**
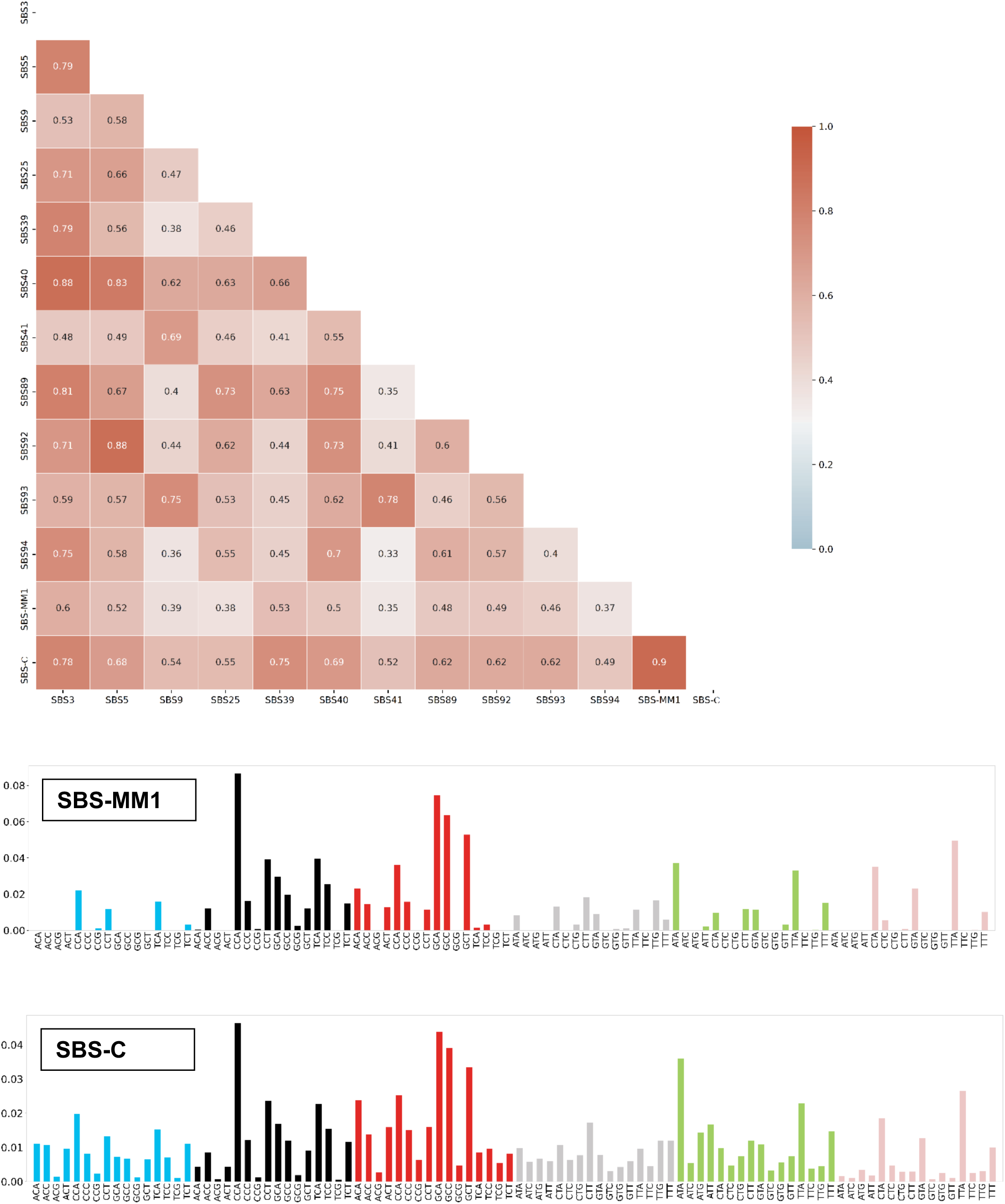
The pairwise association plot displays the cosine similarity metrics between pairs of mutation signatures, with the strength of associations represented by color intensity. The actual mutational signature plots for SBS-MM1 and SBS-C are also shown.

**Extended Figure 5.**
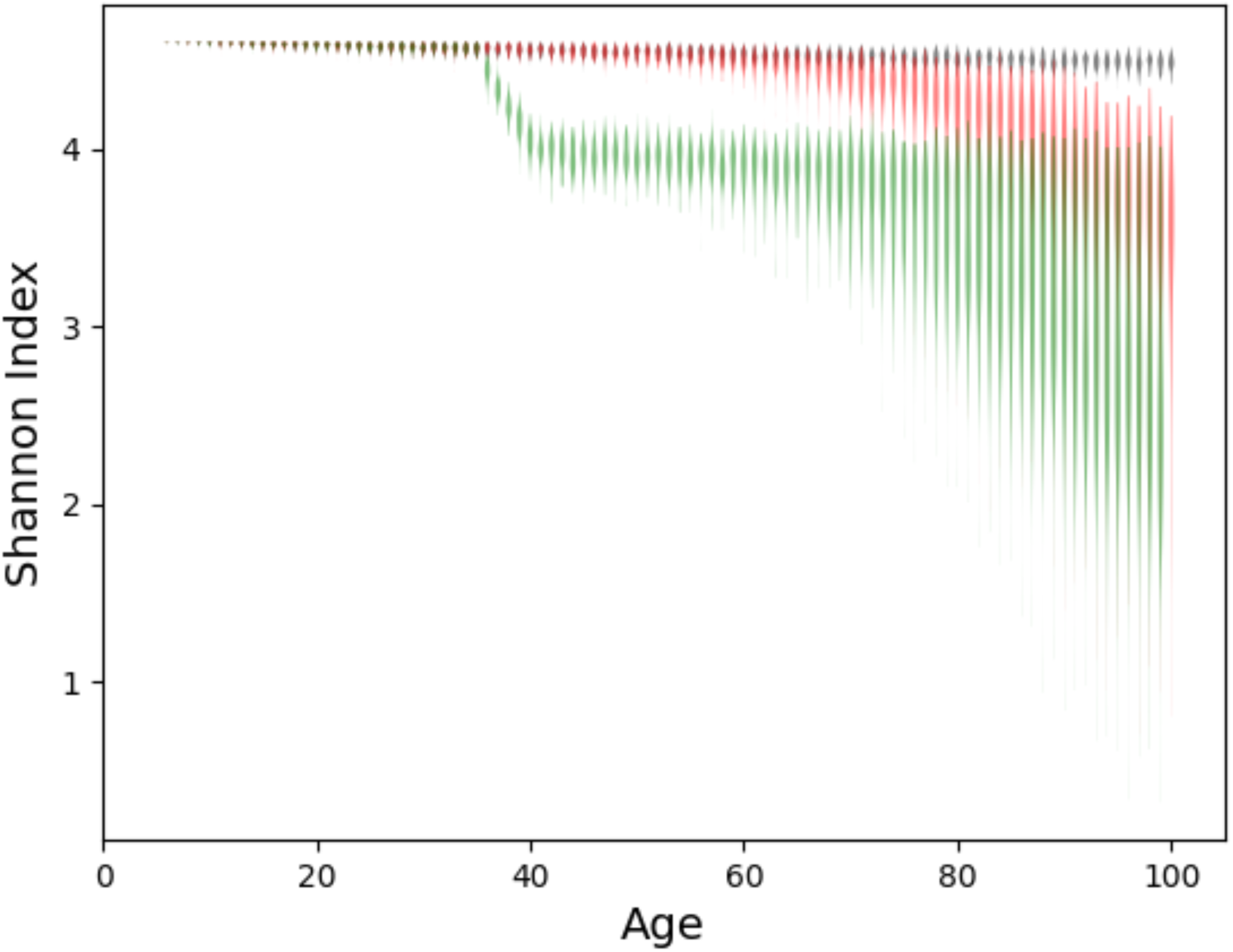
Simulation of Shannon diversity index over time when 100 HSPCs are randomly sampled from 100,000 HSPC pool. Black line assumes that acquired mutations have neutral fitness. Red line assumes that some mutations have selective advantage. Green line assumes that chemotherapy is administered around age 35 to 40.

**Extended Figure 6.**
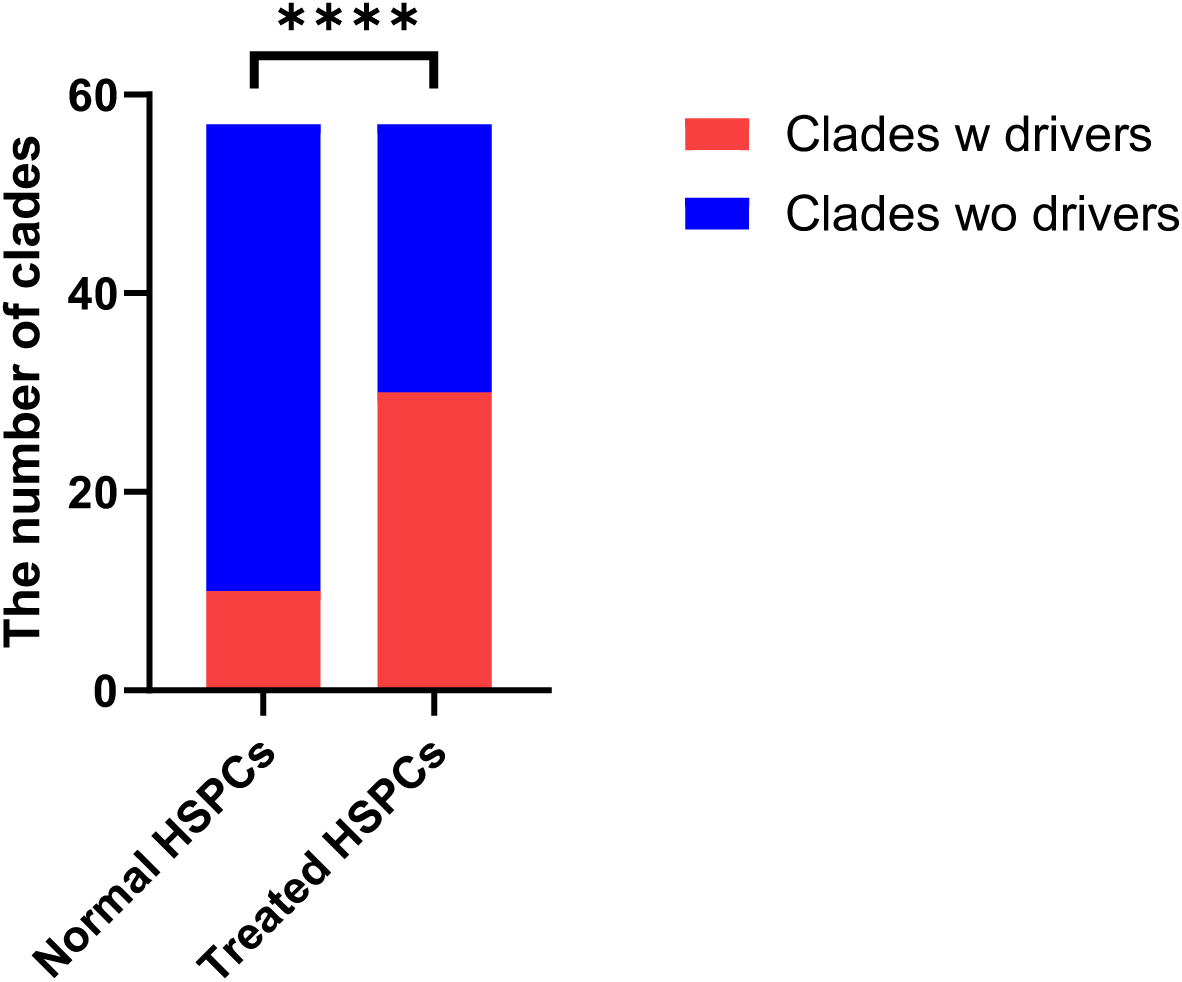
Comparative analysis of clades harboring driver mutations in normal and treated hematopoietic stem and progenitor cells (HSPCs). The graph contrasts the proportion of clades with driver mutations identified in a previously published dataset by Mitchell et al. against those found in the current study’s chemotherapy-treated HSPCs. The statistical significance of the difference is denoted by ****, representing a P-value of less than 0.0001, as determined by the Chi-Square test.

**Extended Figure 7.**
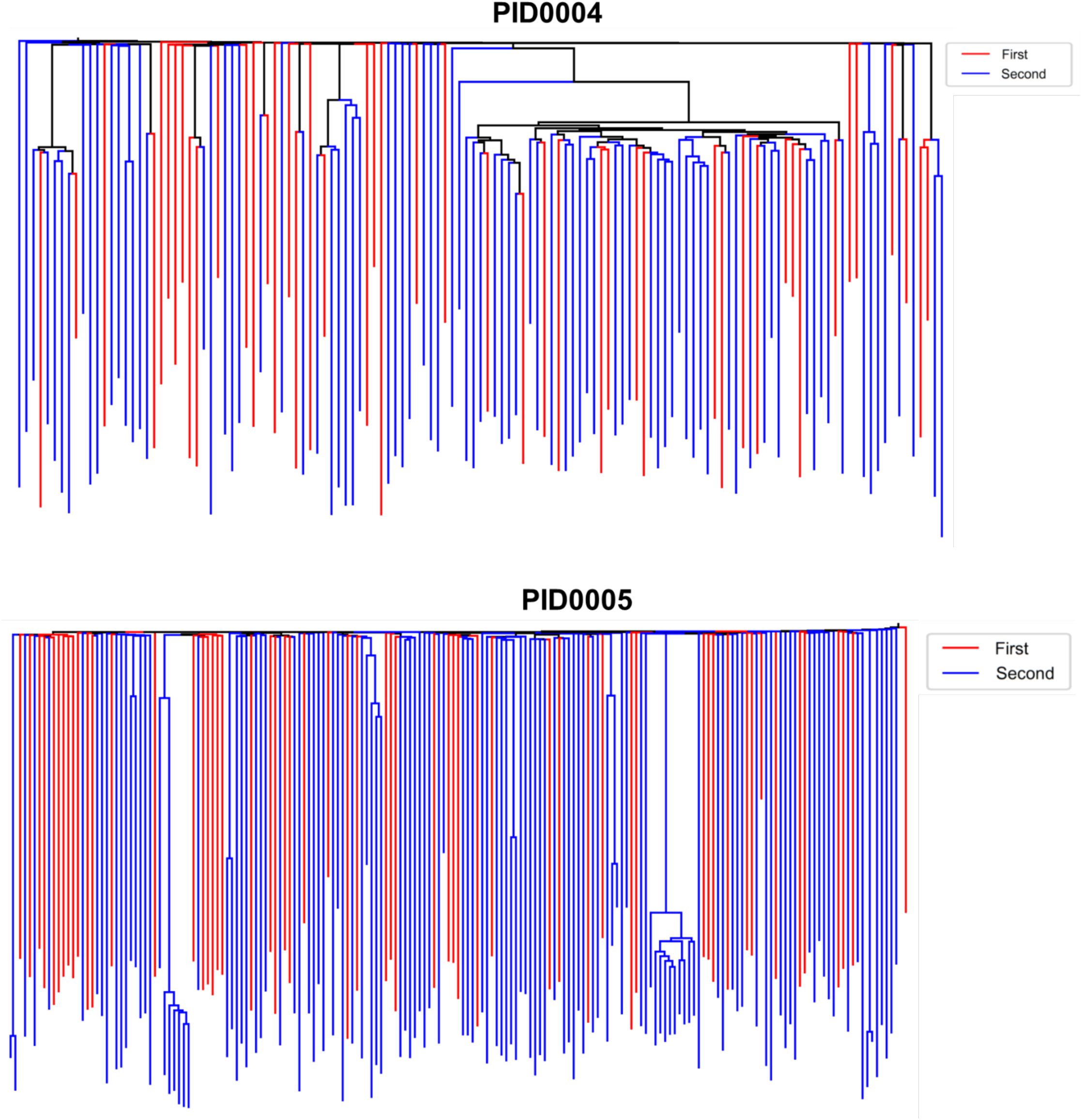
Temporal phylogenetic analysis of HSPC colonies from PID0004 and PID0005. This figure depicts the phylogenetic trees integrating colonies sampled at two different timepoints. For PID0004, the two timepoints are separated by an interval of 3 years, while for PID0005 the interval is 15 years. Colonies collected at the first timepoint are represented in red, while those from the second timepoint are denoted in blue, illustrating the clonal evolution over time.

**Extended Figure 8.**
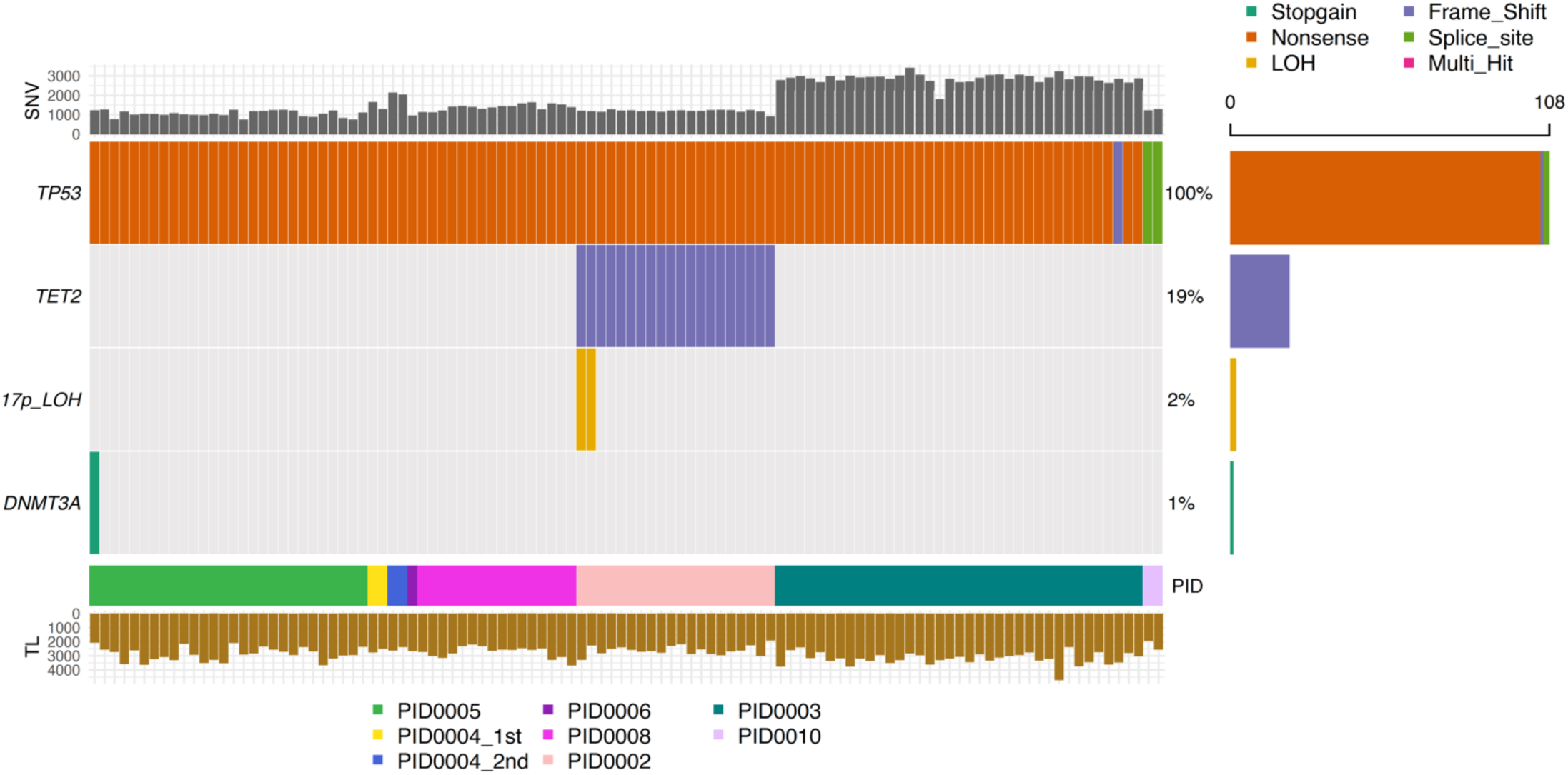
Oncoplot depicting the mutation landscape of 108 HSPC colonies harboring *TP53* mutations. Accompanying driver mutations and copy number alterations are also displayed. The upper bar graph enumerates the total single nucleotide variants (SNVs) identified in each colony, while the lower bar graph presents the estimated telomere lengths across these colonies.

**Extended Figure 9.**
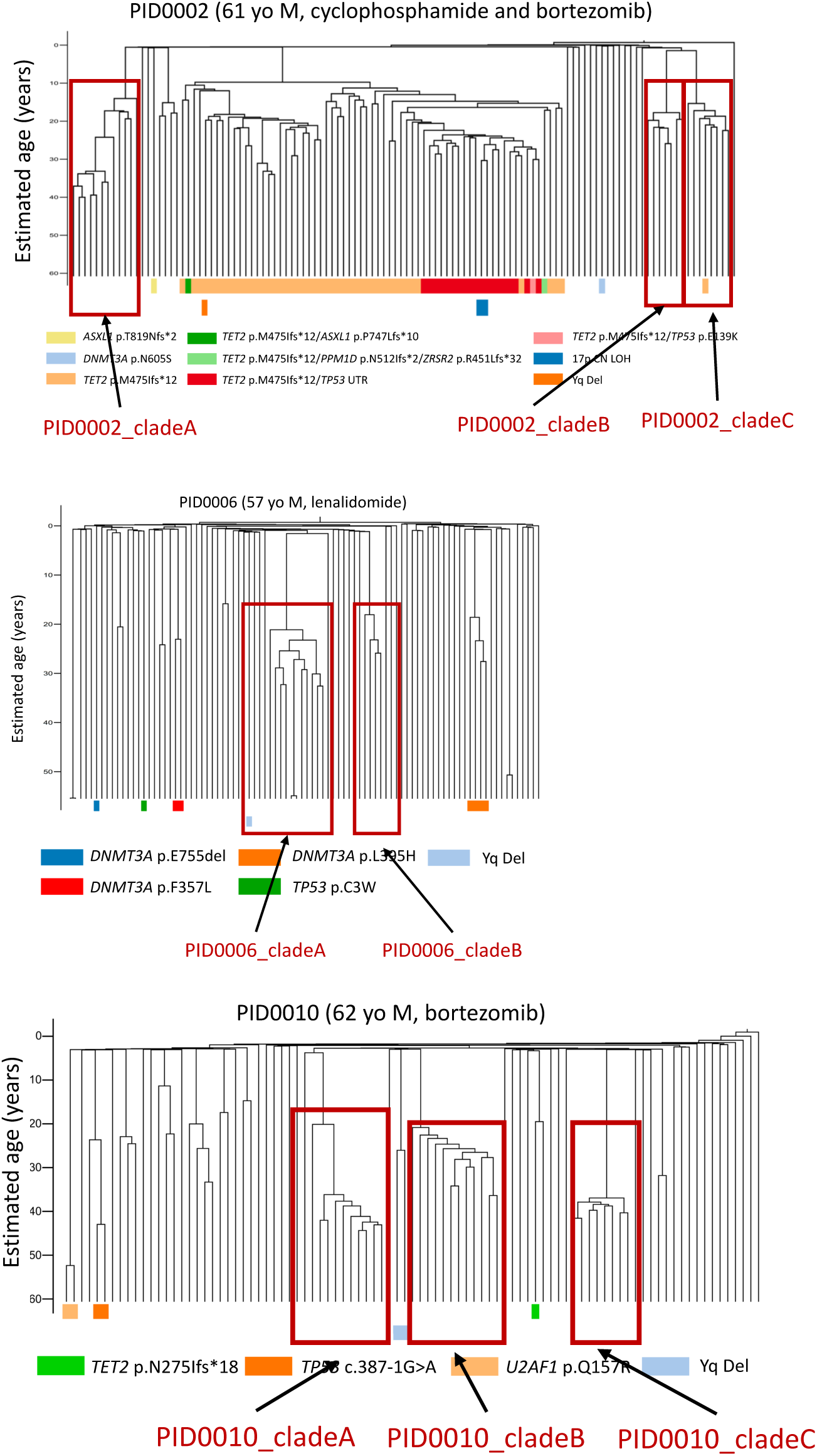
The definition of clades without obvious driver mutations. These clades are used in the analysis of **Figure 5c** and **Figure 5d**.

**Extended Figure 10.**
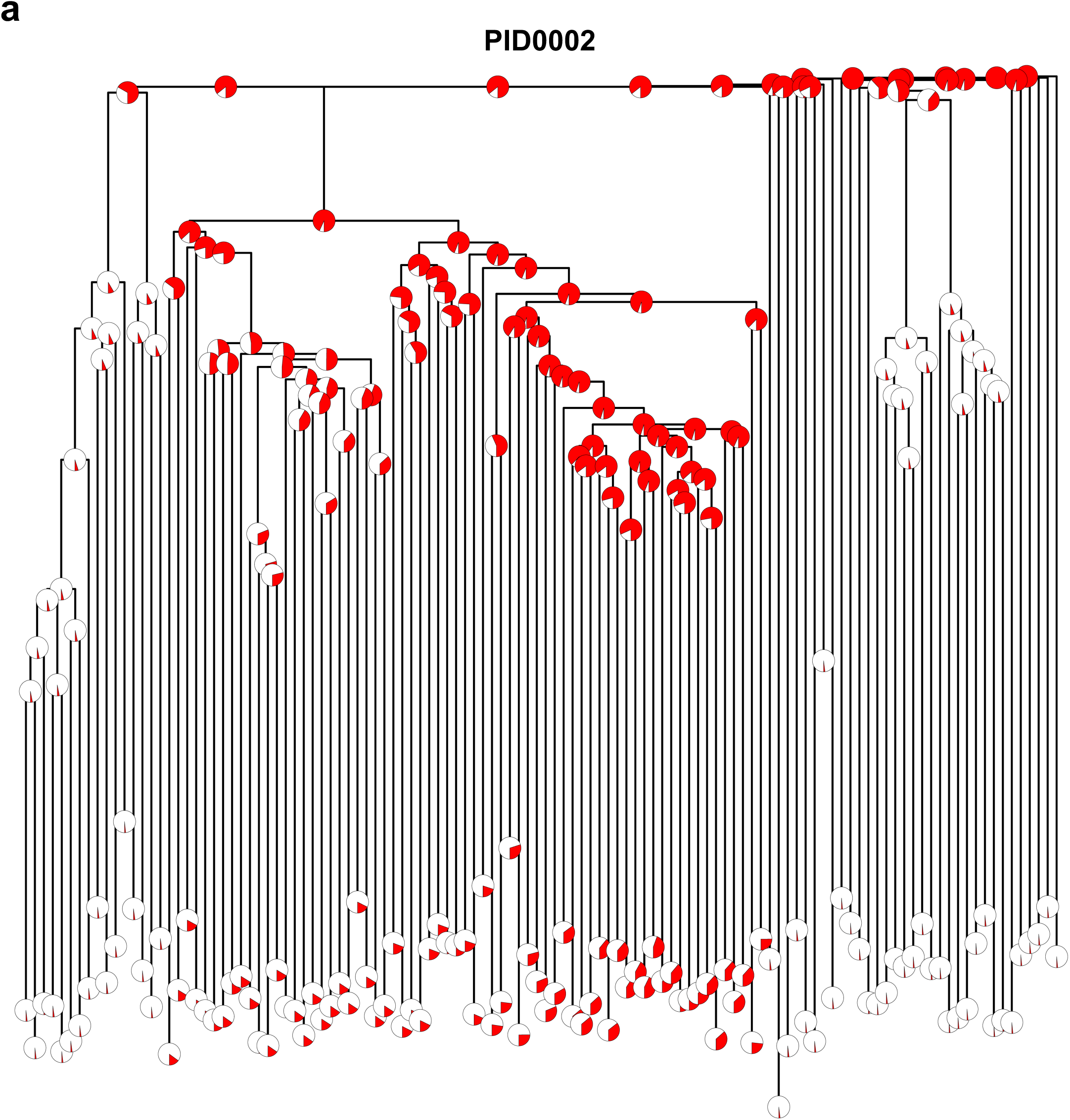
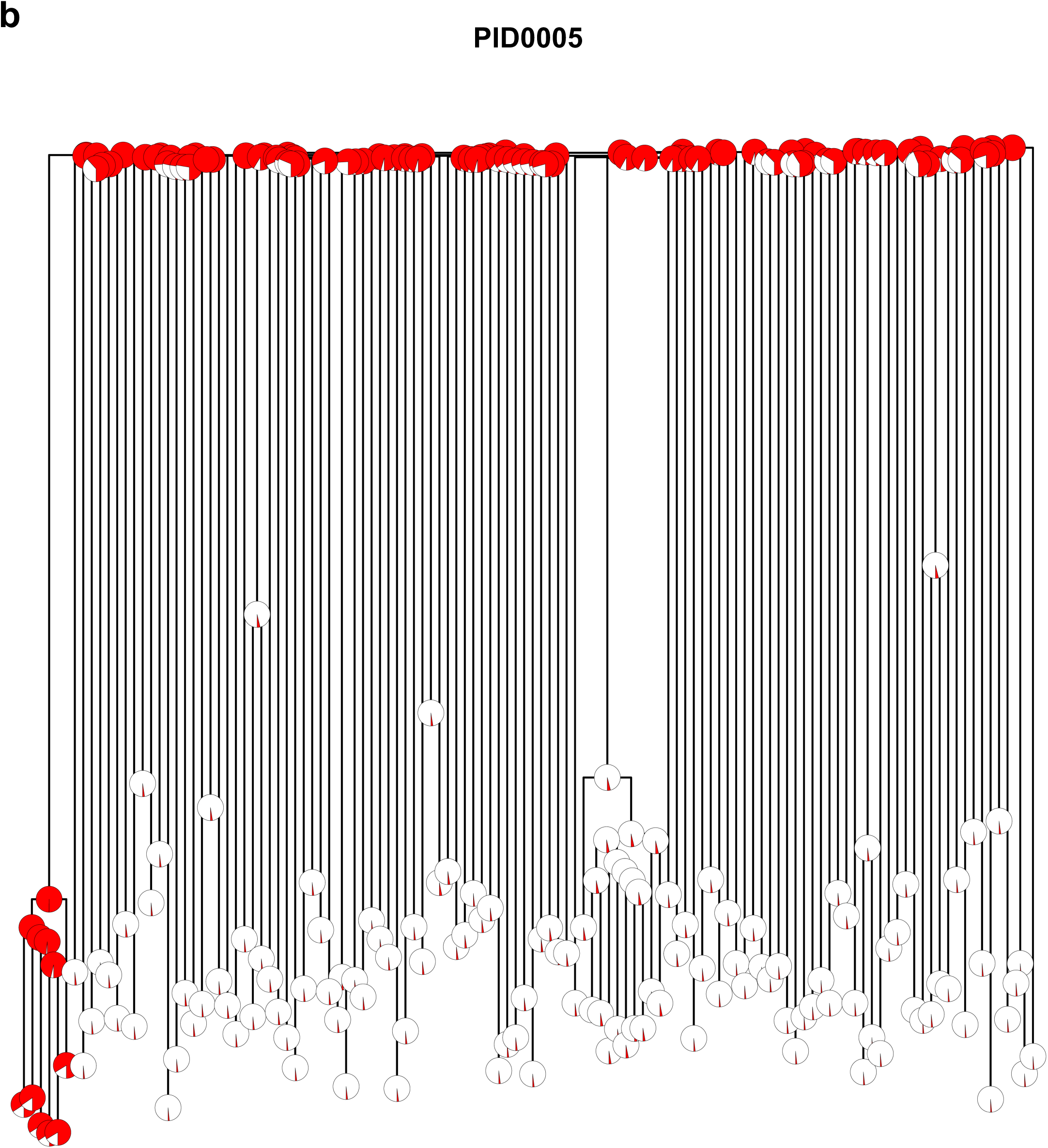
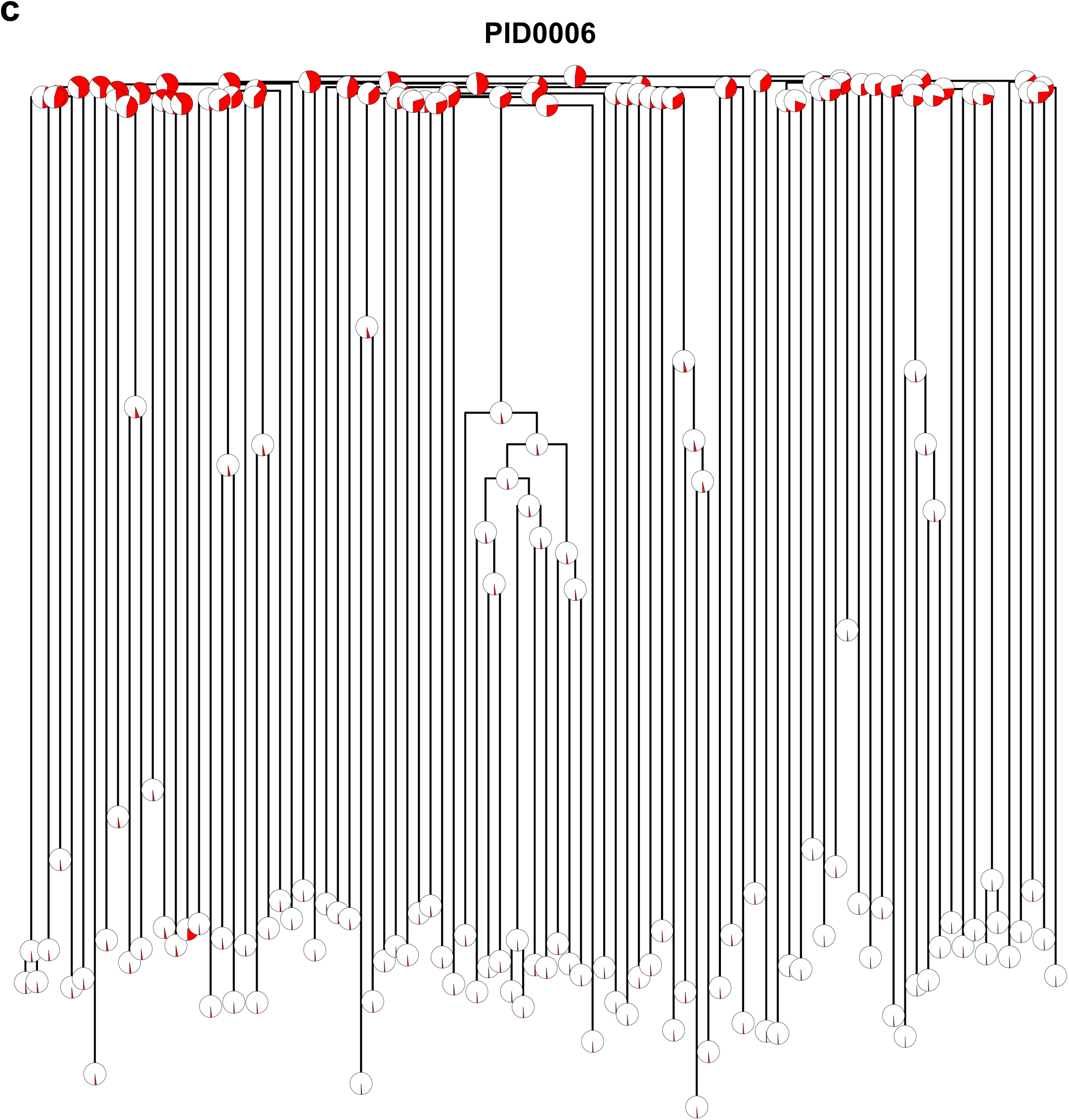
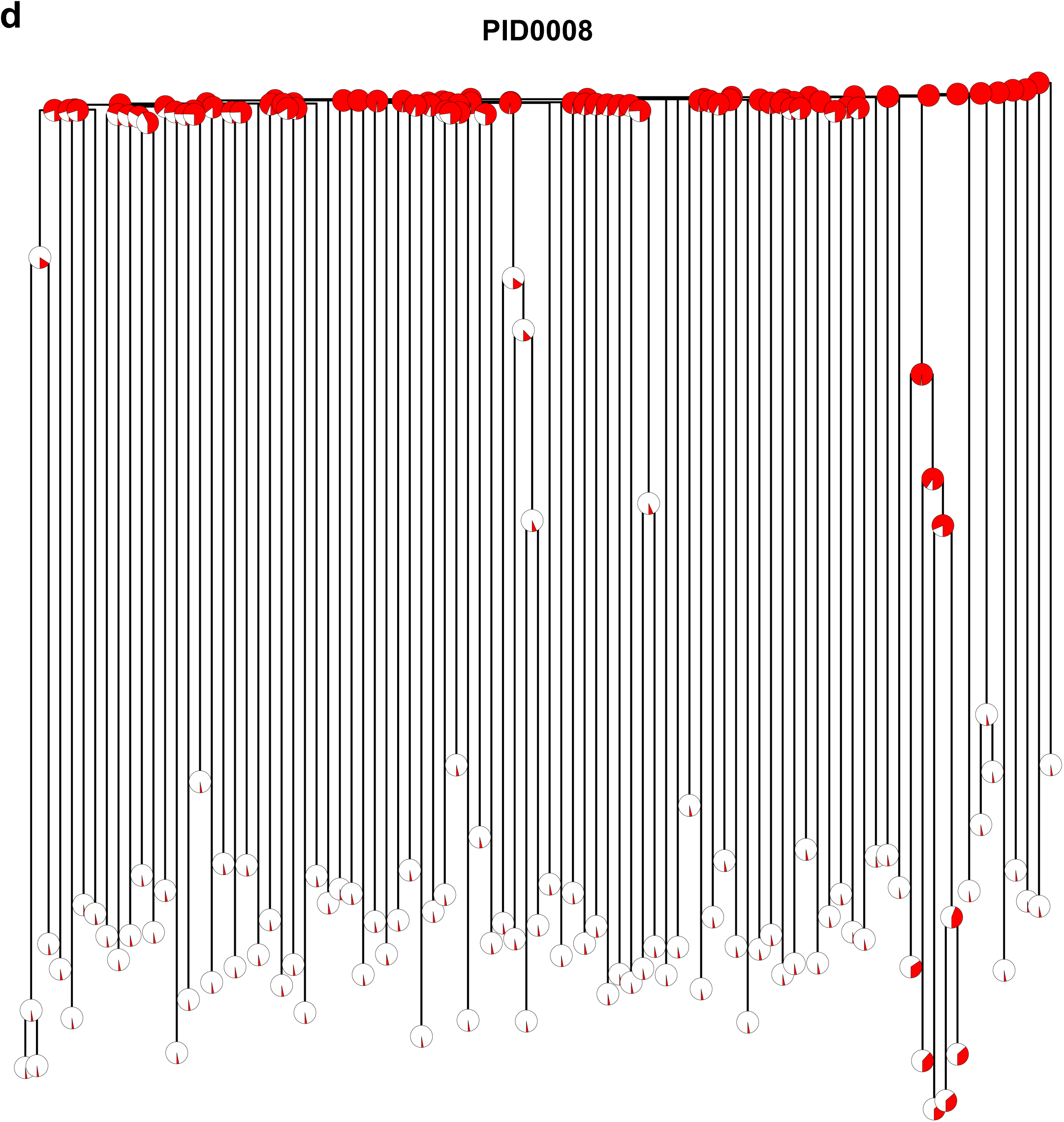
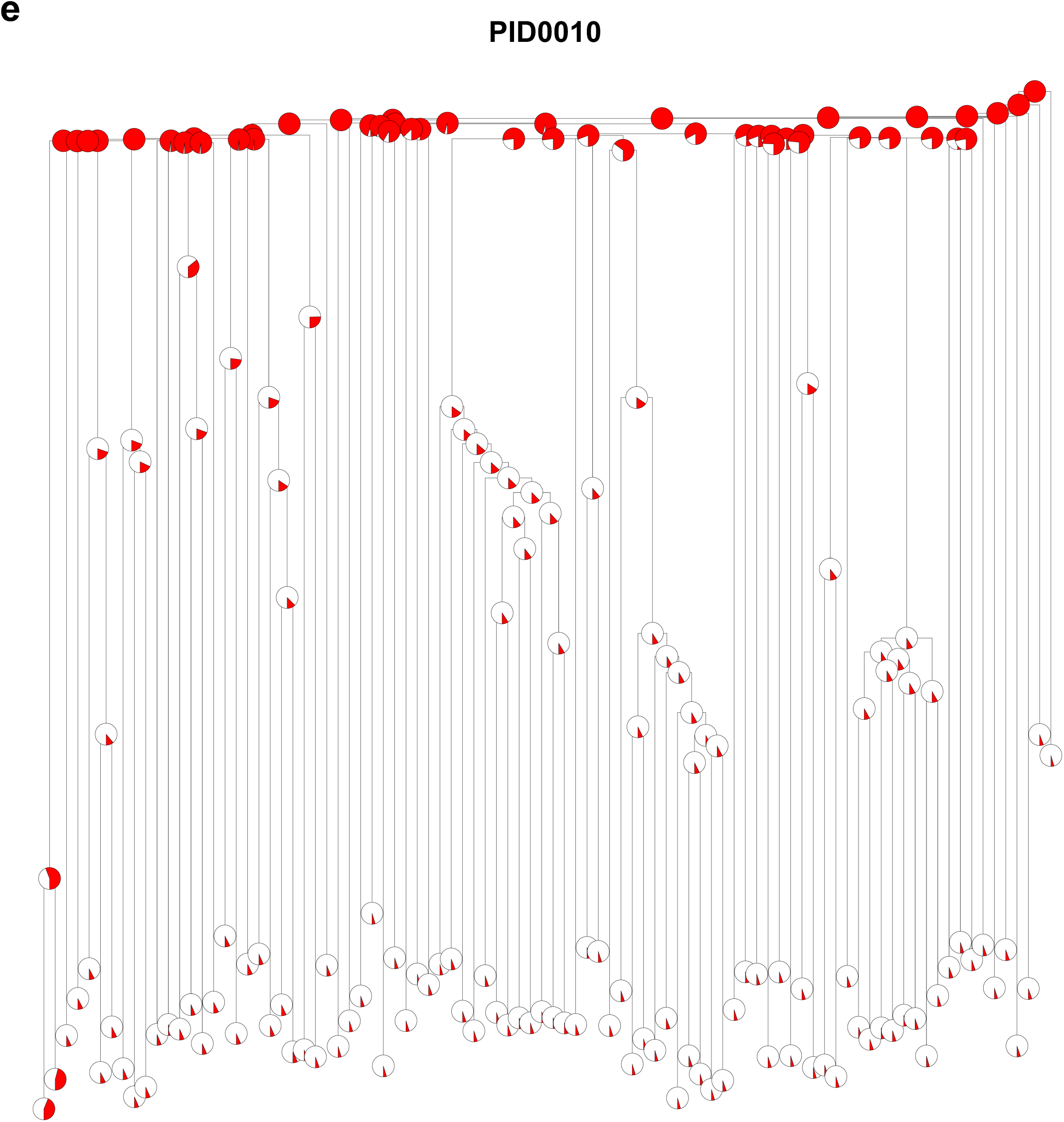
Phylogenetic trees integrating HSPC colonies and matched t-MN genomes in samples where MRCA was identified. The proportion of shared variants between individual colonies and t-MN samples are shown by the pie chart layered onto the trees. (**a**) PID0002, (**b**) PID0005, (**c**) PID0006, (**d**) PID0008, and (**e**) PID0010.

**Extended Figure 11.**
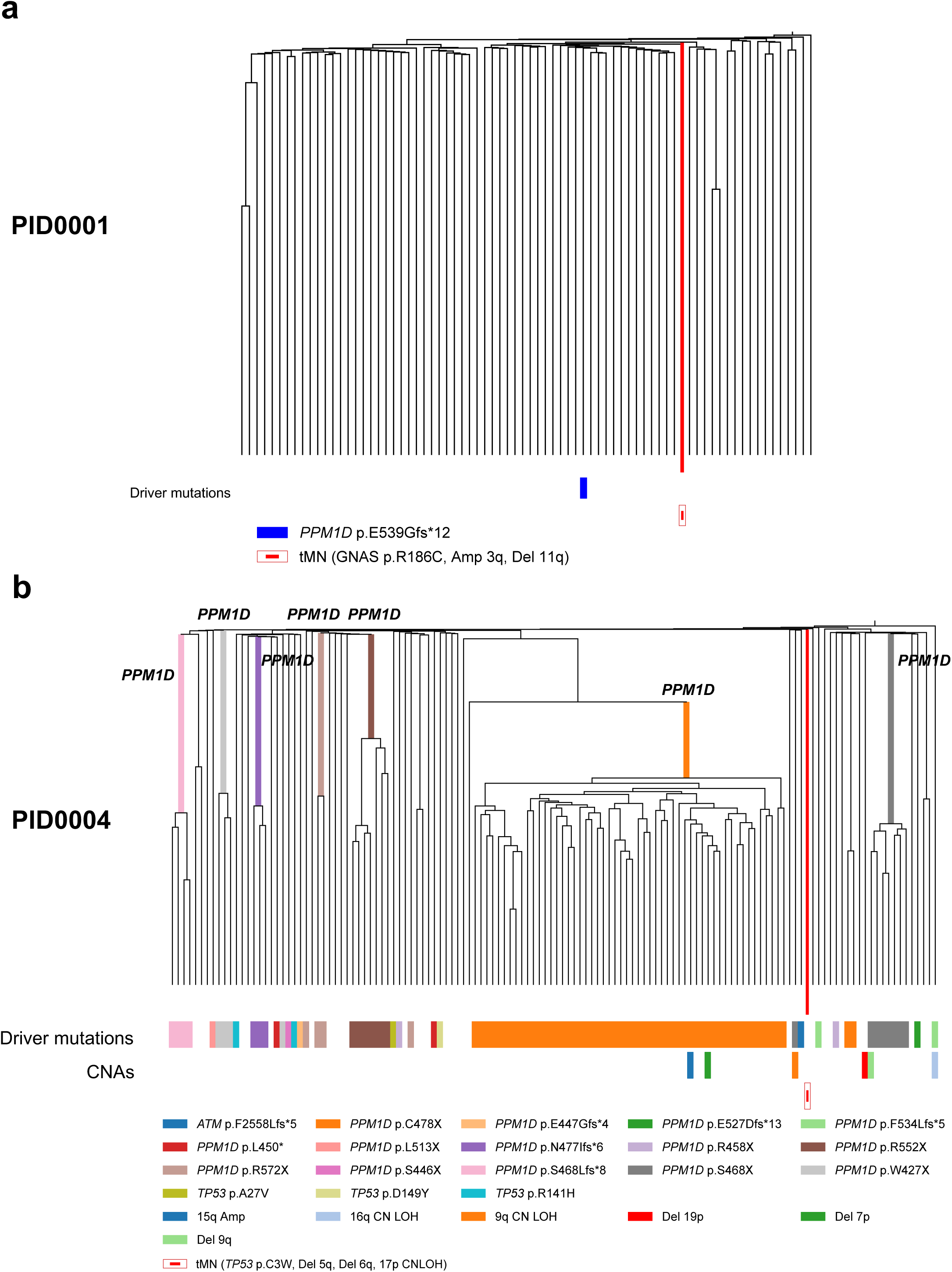
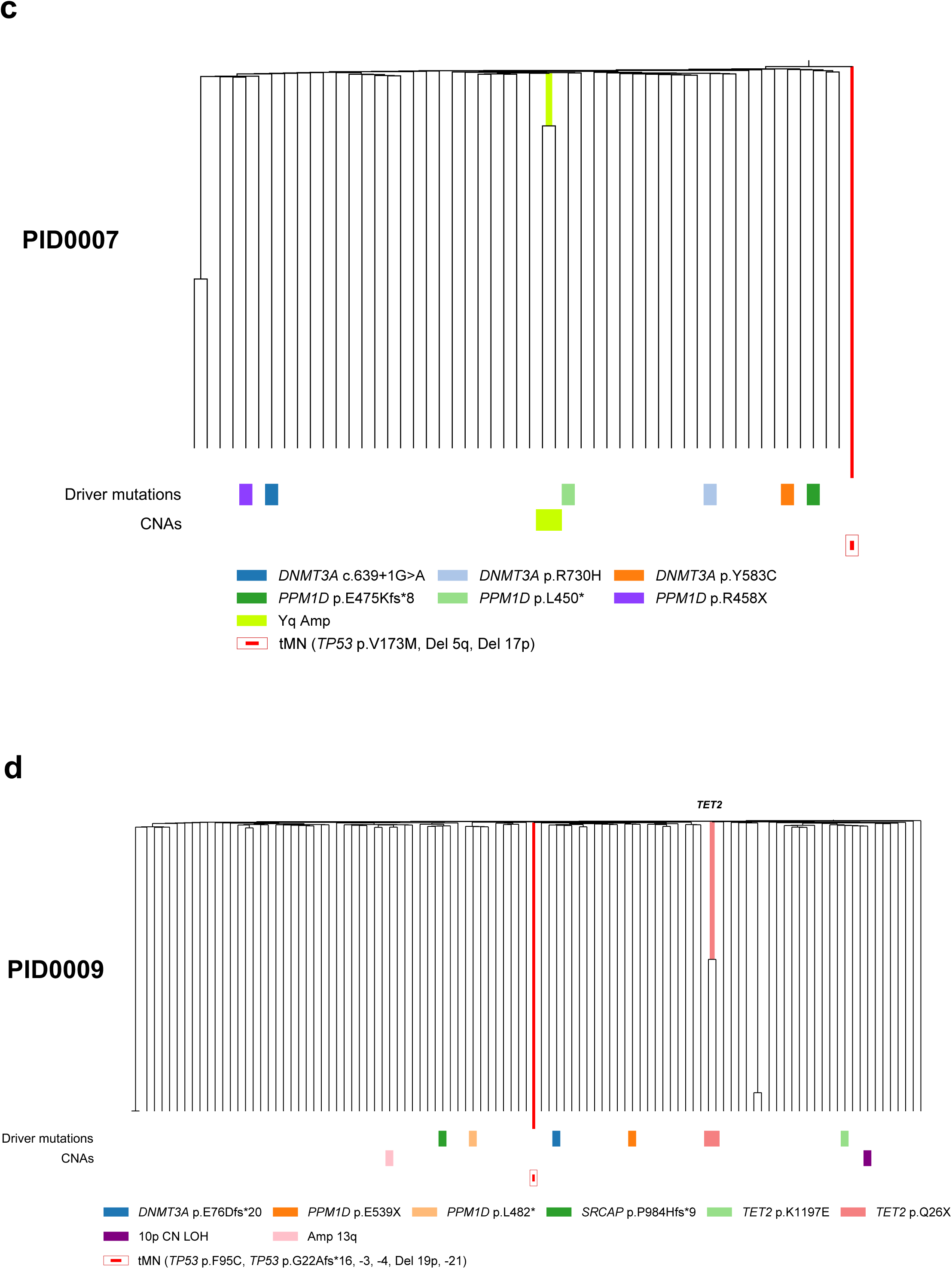
Integrated phylogenetic trees illustrating the clonal evolution of post-treatment HSPCs and corresponding t-MN samples, with a focus on cases where the most recent clonal ancestor (MRCA) could not be determined. The phylogenetic tree for (**a**) PID0001(**b**) PID0004 (**c**) PID0007, and (**d**) PID0009, all displaying the clonal structure without a discernible MRCA.

**Extended Figure 12.**
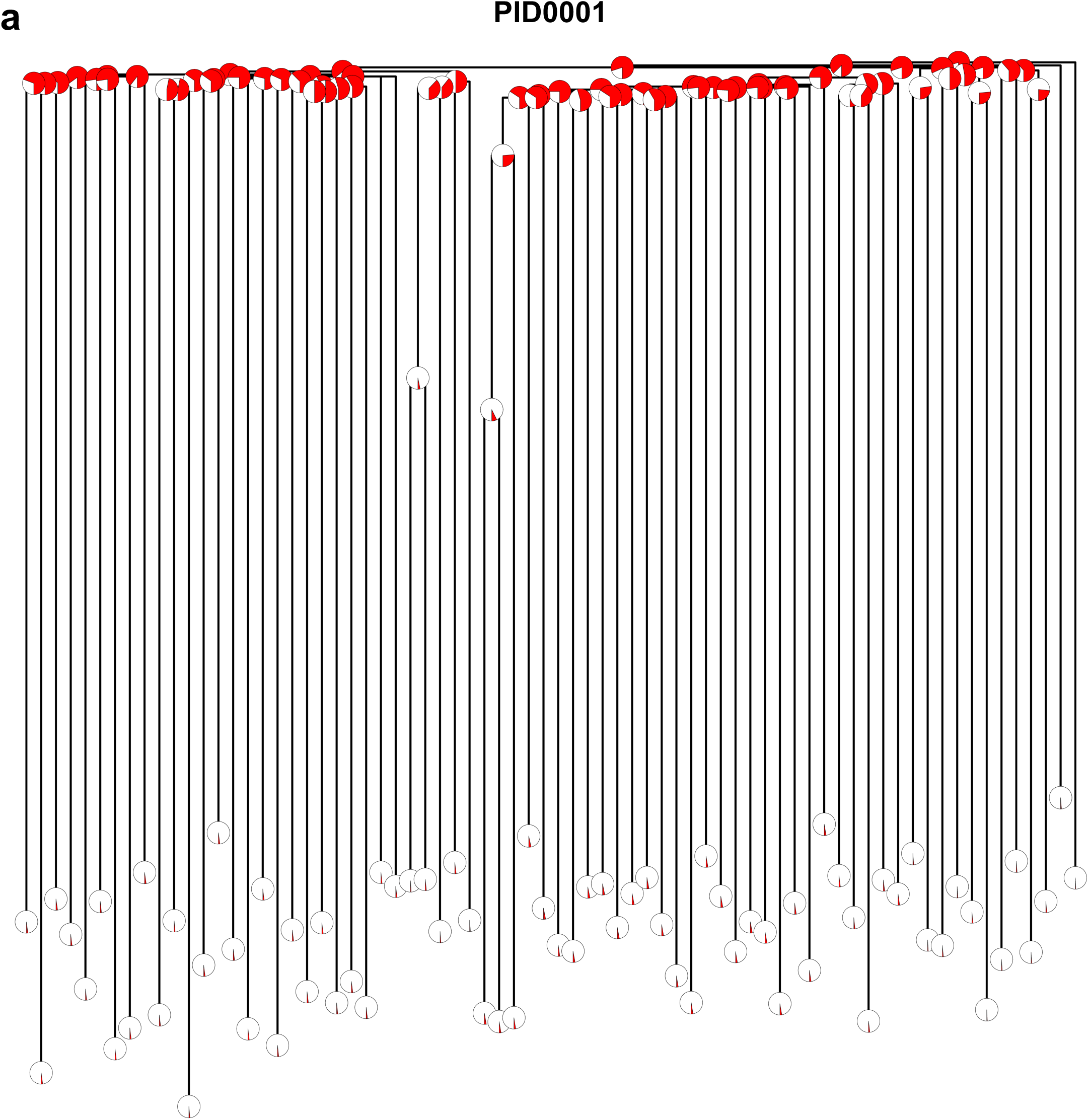
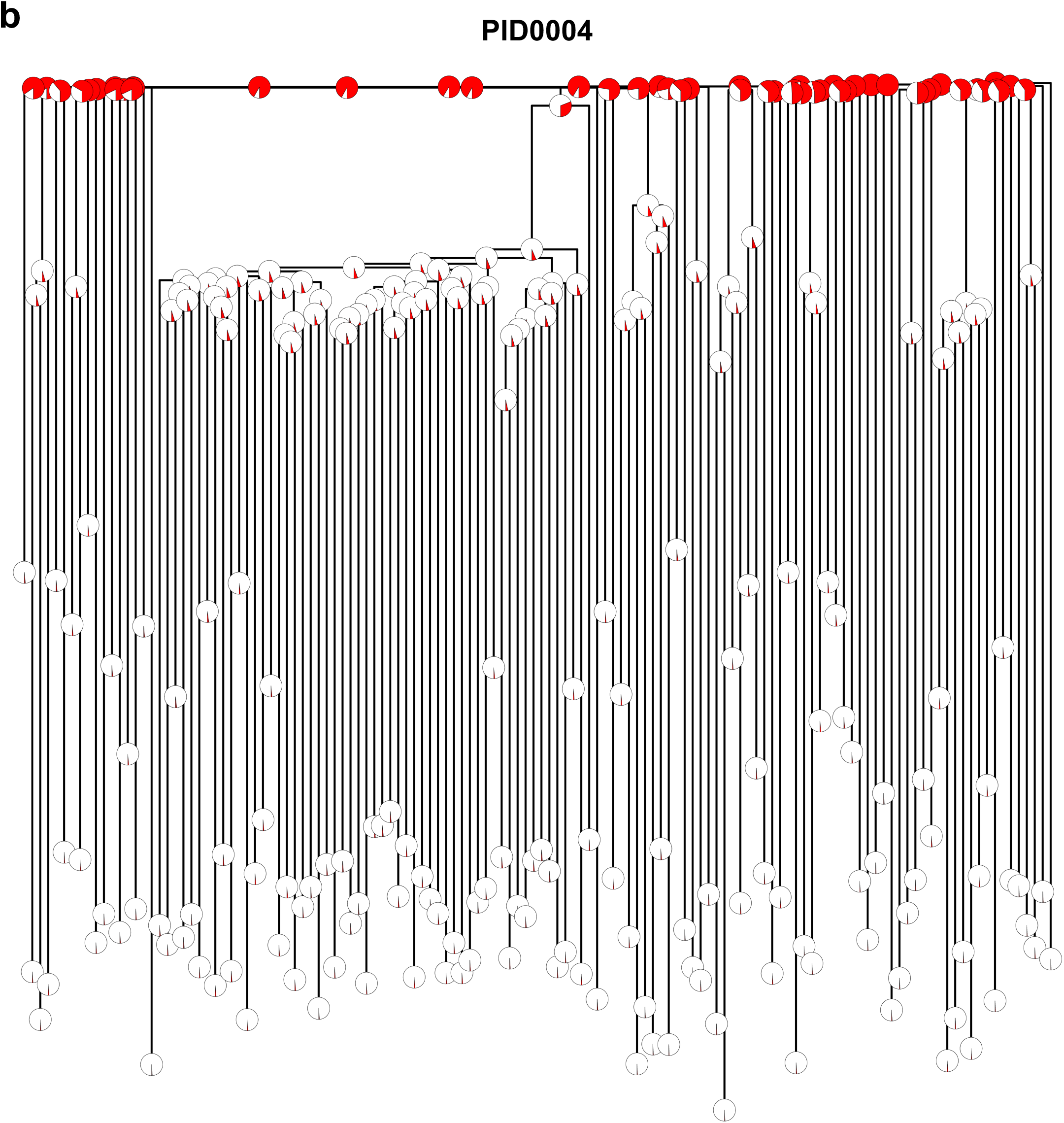
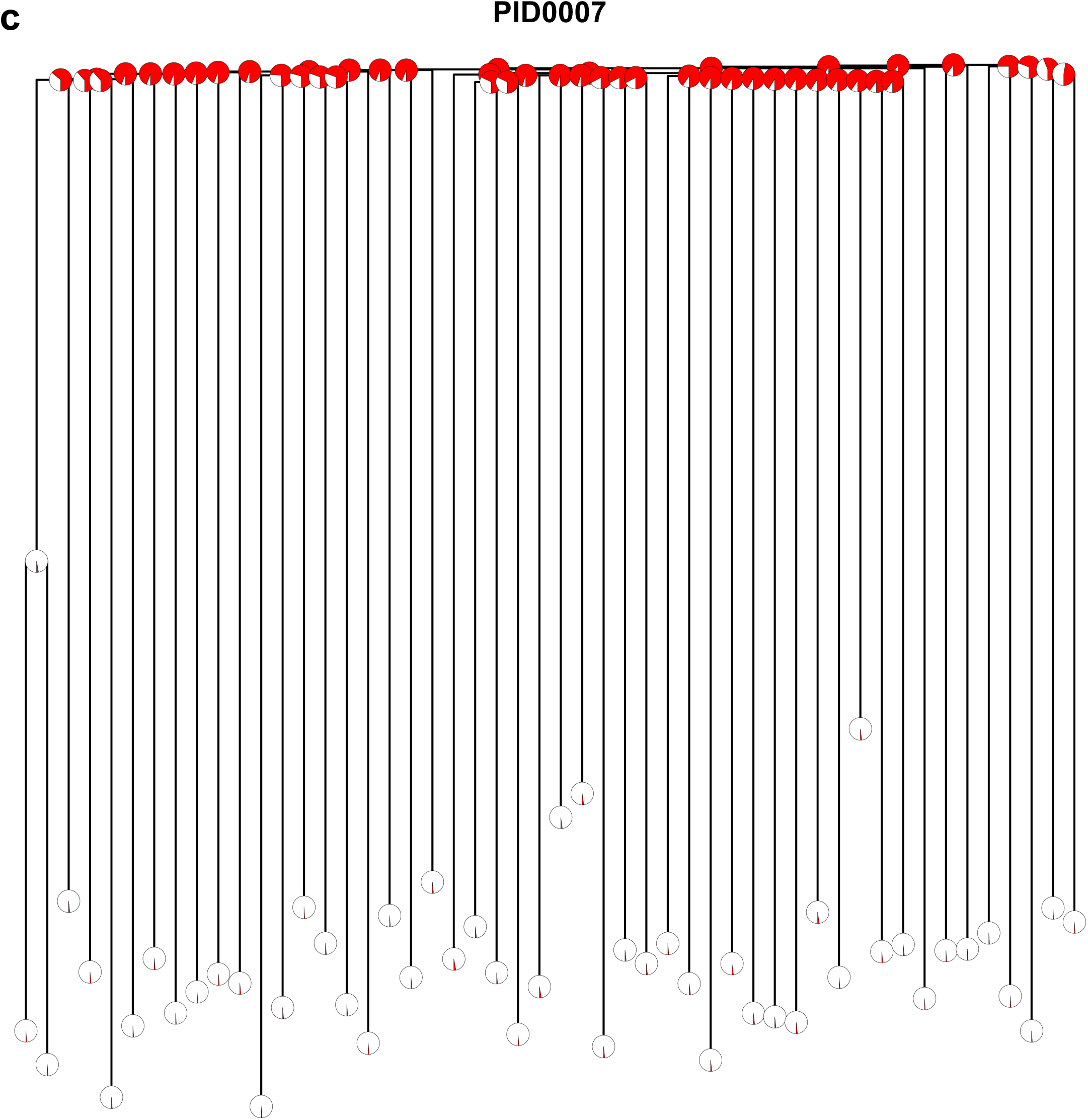
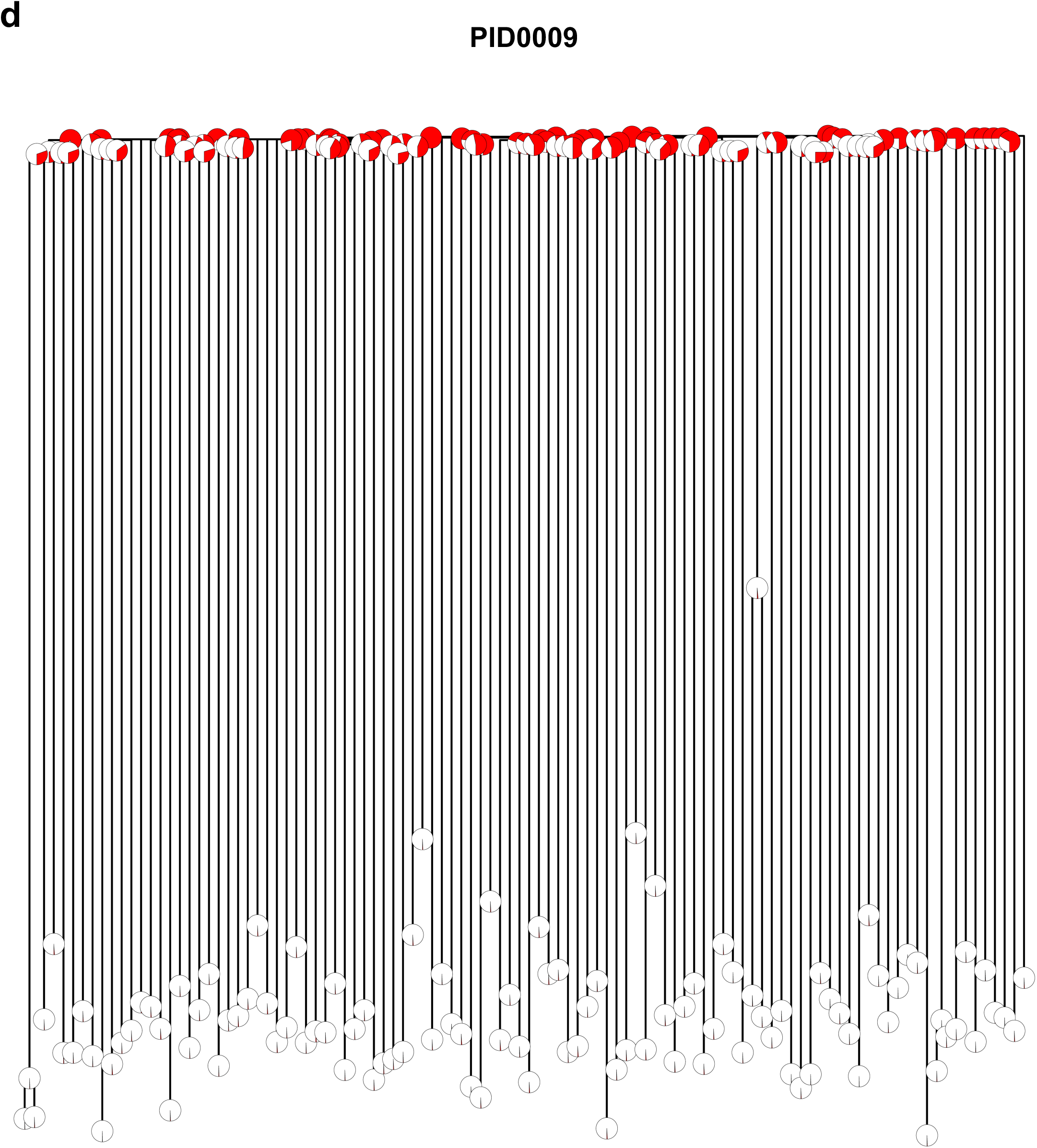
Phylogenetic trees integrating HSPC colonies and matched t-MN genomes in samples where MRCA was not identified. The proportion of shared variants between individual colonies and t-MN samples are shown by the pie chart layered onto the trees. (**a**) PID0001 (**b**) PID0004, (**c**) PID0007, and (**d**) PID0009

**Extended Figure 13.**
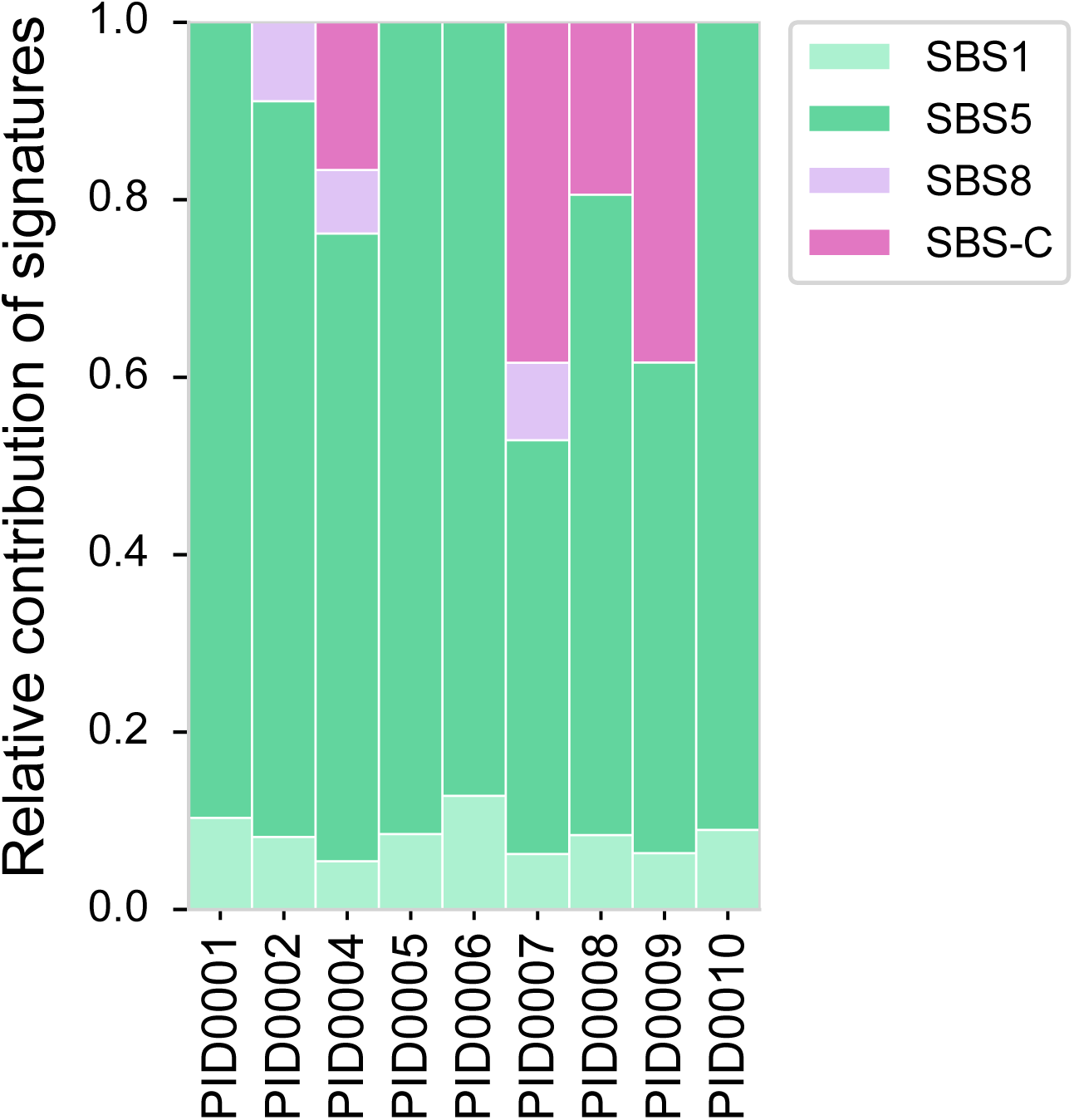
Relative contribution of the four mutational signatures in individual t-MN samples. SBS-C was detected in PID0004, PID0007, PID0008, and PID0009. PID0004 received melphalan during induction therapy. PID0008 bone marrow contained myeloma cells, therefore, SBS-C might be derived from myeloma cells treated with melphalan. For PID0007 and PID0009, the exposure to melphalan only occurred during the conditioning therapy, suggesting that t-MN might have arisen from residual bone marrow cells exposed to melphalan conditioning. PID0008 also contained SBS-C, however, the sample also contained persistent myeloma cells, making it difficult to discern whether SBS-C is from t-MN genome or myeloma cell genome.

**Extended Figure 14.**
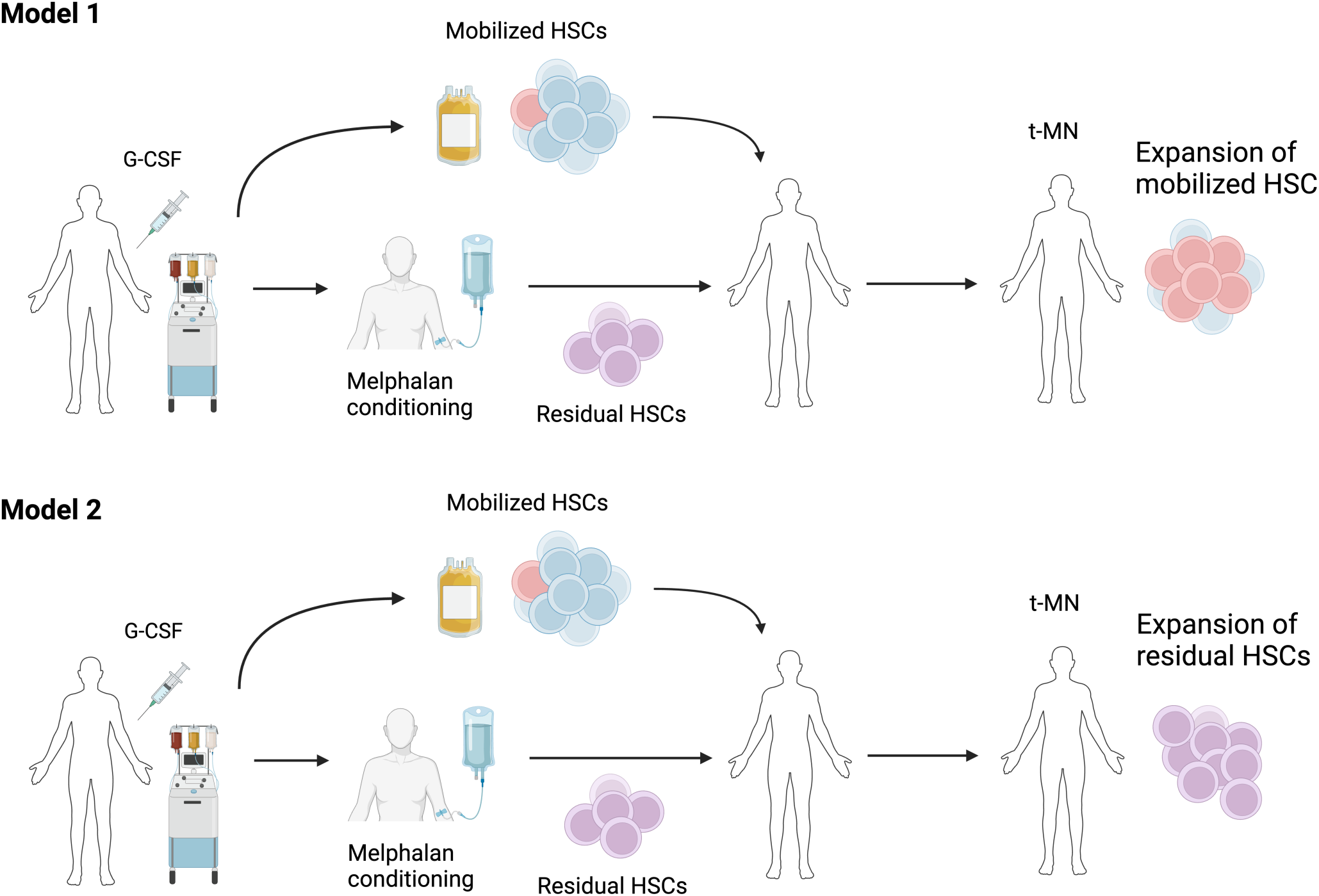
Illustration of two hypothesized models for the development of therapy-related myeloid neoplasms (t-MNs). Model 1 depicts the scenario where t-MNs originate from hematopoietic stem cells (HSCs) that have been mobilized and subsequently transplanted. Model 2 represents the alternative pathway where t-MNs develop from residual HSCs remaining in the bone marrow, which were not mobilized and thus are subject to high-dose melphalan conditioning regimens.

## References

1 Mitchell, E. et al. Clonal dynamics of haematopoiesis across the human lifespan. Nature 606, 343–350, doi:10.1038/s41586-022-04786-y (2022).

2 Vijg, J. & Dong, X. Pathogenic Mechanisms of Somatic Mutation and Genome Mosaicism in Aging. Cell 182, 12–23, doi:10.1016/j.cell.2020.06.024 (2020).

3 Abascal, F. et al. Somatic mutation landscapes at single-molecule resolution. Nature 593, 405–410, doi:10.1038/s41586-021-03477-4 (2021).

4 Lee-Six, H. et al. Population dynamics of normal human blood inferred from somatic mutations. Nature, doi:10.1038/s41586-018-0497-0 (2018).

5 Welch, J. S. et al. The origin and evolution of mutations in acute myeloid leukemia. Cell 150, 264–278, doi:10.1016/j.cell.2012.06.023 (2012).

6 Jaiswal, S. et al. Age-related clonal hematopoiesis associated with adverse outcomes. N Engl J Med 371, 2488–2498, doi:10.1056/NEJMoa1408617 (2014).

7 Genovese, G. et al. Clonal hematopoiesis and blood-cancer risk inferred from blood DNA sequence. N Engl J Med 371, 2477–2487, doi:10.1056/NEJMoa1409405 (2014).

8 Fabre, M. A. et al. The longitudinal dynamics and natural history of clonal haematopoiesis. Nature 606, 335–342, doi:10.1038/s41586-022-04785-z (2022).

9 Martincorena, I. et al. High burden and pervasive positive selection of somatic mutations in normal human skin. Science 348, 880–886, doi:doi:10.1126/science.aaa6806 (2015).

10 Yoshida, K. et al. Tobacco smoking and somatic mutations in human bronchial epithelium. Nature 578, 266–272, doi:10.1038/s41586-020-1961-1 (2020).

11 Huang, Z. et al. Single-cell analysis of somatic mutations in human bronchial epithelial cells in relation to aging and smoking. Nat Genet 54, 492–498, doi:10.1038/s41588-022-01035-w (2022).

12 Yokoyama, A. et al. Age-related remodelling of oesophageal epithelia by mutated cancer drivers. Nature, 1, doi:doi:10.1038/s41586-018-0811-x (2019).

13 Martincorena, I. et al. Somatic mutant clones colonize the human esophagus with age. Science 362, 911–917, doi:doi:10.1126/science.aau3879 (2018).

14 Hill, W. et al. Lung adenocarcinoma promotion by air pollutants. Nature 616, 159–167, doi:10.1038/s41586-023-05874-3 (2023).

15 McNerney, M. E., Godley, L. A. & Le Beau, M. M. Therapy-related myeloid neoplasms: when genetics and environment collide. Nat Rev Cancer 17, 513–527, doi:10.1038/nrc.2017.60 (2017).

16 Coombs, C. C. et al. Therapy-related clonal hematopoiesis in patients with non-hematologic cancers is common and associated with adverse clinical outcomes. Cell stem cell 21, 374–382. e374 (2017).

17 Bolton, K. L. et al. Cancer therapy shapes the fitness landscape of clonal hematopoiesis. Nat Genet 52, 1219–1226, doi:10.1038/s41588-020-00710-0 (2020).

18 Hsu, J. I. et al. PPM1D Mutations Drive Clonal Hematopoiesis in Response to Cytotoxic Chemotherapy. Cell Stem Cell 23, 700–713 e706, doi:10.1016/j.stem.2018.10.004 (2018).

19 Wong, T. N. et al. Role of TP53 mutations in the origin and evolution of therapy-related acute myeloid leukaemia. Nature 518, 552–555, doi:10.1038/nature13968 (2015).

20 Sperling, A. S. et al. Lenalidomide promotes the development of TP53-mutated therapy-related myeloid neoplasms. Blood 140, 1753–1763, doi:10.1182/blood.2021014956 (2022).

21 Rustad, E. H. et al. Timing the initiation of multiple myeloma. Nature Communications 11, 1917, doi:10.1038/s41467-020-15740-9 (2020).

22 Brunton, L. L. & Knollmann, B. r. C. 1 online resource (McGraw Hill, New York, 2023).

23 Alexandrov, L. B. et al. Clock-like mutational processes in human somatic cells. Nat Genet 47, 1402–1407, doi:10.1038/ng.3441 (2015).

24 Hess, D. A. et al. Selection based on CD133 and high aldehyde dehydrogenase activity isolates long-term reconstituting human hematopoietic stem cells. Blood 107, 2162–2169, doi:10.1182/blood-2005-06-2284 (2006).

25 Emadi, A., Jones, R. J. & Brodsky, R. A. Cyclophosphamide and cancer: golden anniversary. Nature Reviews Clinical Oncology 6, 638–647, doi:10.1038/nrclinonc.2009.146 (2009).

26 Moran, P. A. P. Random processes in genetics. Mathematical Proceedings of the Cambridge Philosophical Society 54, 60–71, doi:10.1017/S0305004100033193 (1958).

27 de Kanter, J. K. et al. Antiviral treatment causes a unique mutational signature in cancers of transplantation recipients. Cell Stem Cell 28, 1726–1739 e1726, doi:10.1016/j.stem.2021.07.012 (2021).

28 Bolaños-Meade, J. et al. Post-Transplantation Cyclophosphamide-Based Graft-versus-Host Disease Prophylaxis. New Engl J Med 388, 2338–2348, doi:10.1056/NEJMoa2215943 (2023).

29 Curtis, R. E. et al. Risk of Leukemia after Chemotherapy and Radiation Treatment for Breast Cancer. New Engl J Med 326, 1745–1751, doi:10.1056/nejm199206253262605 (1992).

30 Takahashi, K. et al. Preleukaemic clonal haemopoiesis and risk of therapy-related myeloid neoplasms: a case-control study. Lancet Oncol 18, 100–111, doi:10.1016/S1470-2045(16)30626-X (2017).

31 Wong, T. N. et al. Role of TP53 mutations in the origin and evolution of therapy-related acute myeloid leukaemia. Nature 518, 552–555, doi:10.1038/nature13968 (2015).

32 Abelson, S. et al. Prediction of acute myeloid leukaemia risk in healthy individuals. Nature 559, 400–404, doi:10.1038/s41586-018-0317-6 (2018).

33 Desai, P. et al. Somatic mutations precede acute myeloid leukemia years before diagnosis. Nat Med 24, 1015–1023, doi:10.1038/s41591-018-0081-z (2018).

34 Bernard, E. et al. Implications of TP53 allelic state for genome stability, clinical presentation and outcomes in myelodysplastic syndromes. Nat Med 26, 1549–1556, doi:10.1038/s41591-020-1008-z (2020).

35 Diamond, B. et al. Tracking the evolution of therapy-related myeloid neoplasms using chemotherapy signatures. Blood 141, 2359–2371, doi:10.1182/blood.2022018244 (2023).

36 Benjamin, D. et al. Calling Somatic SNVs and Indels with Mutect2. bioRxiv, 861054, doi:10.1101/861054 (2019).

37 Farmery, J. H. R., Smith, M. L., Diseases, N. B.-R. & Lynch, A. G. Telomerecat: A ploidy-agnostic method for estimating telomere length from whole genome sequencing data. Sci Rep 8, 1300, doi:10.1038/s41598-017-14403-y (2018).

38 Alexandrov, L. B. et al. The repertoire of mutational signatures in human cancer. Nature 578, 94–101, doi:10.1038/s41586-020-1943-3 (2020).

39 Islam, S. M. A. et al. Uncovering novel mutational signatures by de novo extraction with SigProfilerExtractor. Cell Genomics 2, 100179, (2022).

40 Rustad, E. H. et al. mmsig: a fitting approach to accurately identify somatic mutational signatures in hematological malignancies. Communications Biology 4, 424, doi:10.1038/s42003-021-01938-0 (2021).

